# A Novel Transgenic Reporter to Study Vertebrate Epigenetics

**DOI:** 10.1101/2025.04.21.649821

**Authors:** Miranda Marvel, Aniket V. Gore, Kiyohito Taimatsu, Daniel Castranova, Gennady Margolin, Allison Goldstein, Avery Angell Swearer, Olivia Carpinello, Joshua Combs, Megan Detels, Leah Greenspan, Addison Parker, Yehyun Abby Kim, Jian Ming Khor, Andrew E. Davis, Madeleine Kenton, John Prevedel, Keith Barnes, James Iben, Tianwei Li, Fabio Faucz, Ryan Dale, Brant M. Weinstein

## Abstract

Epigenetic reprogramming contributes to the generation of cellular diversity during vertebrate development but the mechanisms directing this are still not well understood. Large-scale genetic screens have been highly successful in identifying epigenetic regulatory genes in invertebrates such as worms and flies, but similar large-scale genetic screens to identify epigenetic regulators have not been carried out in vertebrates. Here we report a newly generated “EpiTag” zebrafish transgenic reporter line that permits easy cellular-level visualization of epigenetic silencing or activation in living animals during development, gametogenesis, and regeneration. We use the EpiTag reporter to carry out an F3 ENU mutagenesis screen for epigenetic silencing or activating mutants, identifying relevant vertebrate tissue-specific epigenetic regulatory genes including a new epigenetic model for metabolic dysfunction-associated fatty liver disease (MAFLD). The EpiTag reporter line represents a powerful new tool for genetic and experimental analysis of tissue-specific epigenetic gene regulation in vertebrates.

**One Sentence Summary:** EpiTag transgenic zebrafish provide a powerful new tool for visualizing and studying epigenetic regulation in living vertebrate animals.

## Main Text

Multicellular organisms contain a diversity of cell types with a wide range of cellular functions. Although the roles of transcription factors and morphogens in establishing different cell types during development have been extensively studied, the contribution of epigenetic mechanisms such as histone modifications and/or DNA methylation toward generating cellular diversity during development, and the identities of the tissue-specific regulators that direct this, are still not well understood. Large-scale genetic screens have been highly successful in identifying epigenetic regulatory genesin invertebrates such as worms and flies (*1-3*), but similar large-scale genetic screens to identify mutants in epigenetic regulators have not yet been carried out in vertebrates, in part due to lack of an *in vivo* reporter for the dynamic epigenetic changes occurring during vertebrate development. Here, we describe the generation of a new transgenic zebrafish *in vivo* reporter line that reliably reports tissue-specific epigenetic changes during development, gametogenesis, and regeneration. We use this “EpiTag” reporter to carry out the first large-scale genetic screen in a vertebrate for epigenetic regulatory mutants, identifying a new epigenetic model for metabolic dysfunction-associated fatty liver disease.

Expression of the germ cell-specific *dazl* gene is normally epigenetically silenced in somatic tissues by a “CpG island” (CGI) sequence in its promoter region that is specifically targeted for DNA methylation (4-7). We found the *dazl* CGI is strongly methylated in mature sperm and oocytes and in somatic tissues of developing 5-day old zebrafish larvae but becomes transiently demethylated at the mid-blastula transition (MBT) to early gastrulation stage at approximately 6 hours post-fertilization (hpf; Fig. S1), a stage in development where epigenetic reprogramming is known to take place. Previous reports have shown that the *dazl* CpG island sequence can also promote silencing of adjacent genes in cultured cells (7), so we cloned the zebrafish *dazl* CpG island immediately upstream of a strong, ubiquitously expressed *Xenopus laevis ef1a* promoter driving expression of GFPd2 (8), a destabilized version of green fluorescent protein (half-life reduced from approximately 24 hours to 2 hours) to generate an “EpiTag” reporter transgene (Fig. 1B). We also prepared a control transgene with pCS2 vector sequences of comparable length cloned in place of the *dazl* CpG island (Fig. 1A).

**Figure 1.**
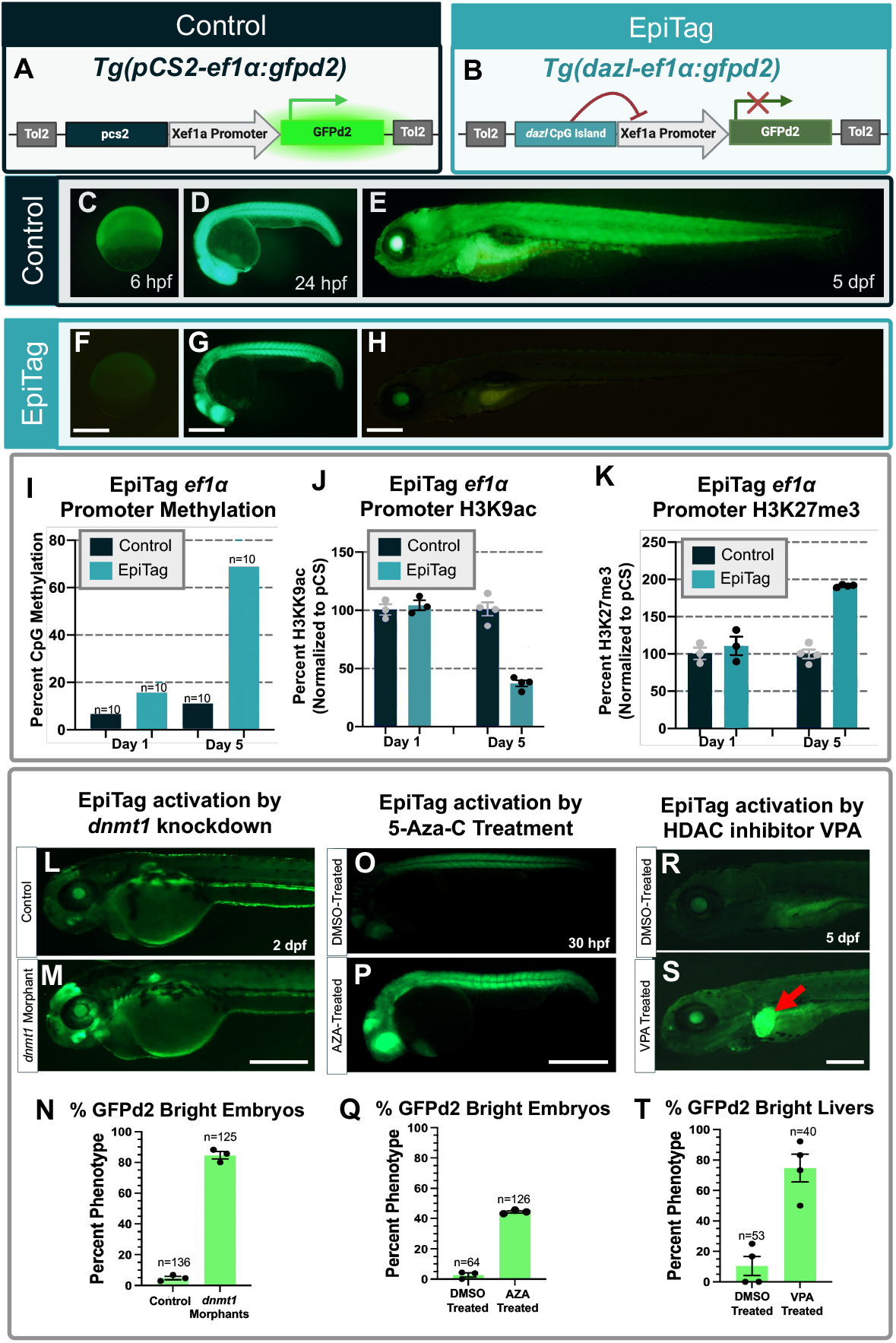
A transgenic “EpiTag” reporter for vertebrate tissue-specific epigenetic silencing. (A,B) Schematic diagram showing the *Tg(pcs-ef1α:gfpd2)*^*y605*^ and *Tg(dazl-ef1α:gfpd2)*^*y603*^ transgenes used to generate stable control and “EpiTag” transgenic lines. (C-H) Lateral view green epifluorescence images of *Tg(pcs-ef1α:gfpd2)*^*y605*^ control (C-E) or *Tg(dazl-ef1α:gfpd2)*^*y603*^ EpiTag (F-H) transgenic animals at 6 hpf (C,F), 24 hpf (D,G) and 5 dpf (E,H). Scale bar, 200 µm. (I-K) Quantification of *Xenopus ef1α* promoter epigenetic modifications in *Tg(pcs-ef1α:gfpd2)*^*y605*^ control (dark navy columns) or *Tg(dazl-ef1α:gfpd2)*^*y603*^ EpiTag (teal columns) animals at 1 dpf or 5 dpf. Graphs show percent CpG methylation measured by targeted bisulfite sequencing (I), and percent H3K9 acetylation (J) or H3K27 trimethylation (K) measured by locus-specific chromatin immunoprecipitation. The values in J and K are normalized to *Tg(pcs-ef1α:gfpd2)*^*y605*^ control levels (set at 100%). (L,M) Lateral view green epifluorescence images of control (L) and *dnmt1* morpholino-injected (M) *Tg(dazl-ef1α:gfpd2)*^*y603*^ EpiTag transgenic animals at 48 hpf. Scale bar, 200 µm. (N) Quantification of the percentage of bright GFPd2 expressing control or *dnmt1* morpholino-injected animals at 48 hpf. (O,P) Green epifluorescence images of control DMSO (O) or 5’-azacytidine (AZA, P) treated *Tg(dazl-ef1α:gfpd2)*^*y603*^ EpiTag transgenic animals at 30 hpf. Scale bar, 200µm. (Q) Quantification of the percentage of bright GFPd2 expressing embryos of control DMSO-treated and AZA-treated EpiTag transgenic animals at 30 hpf. (R,S) Lateral view green epifluorescence images of control DMSO (R) or valproic acid (VPA; S) treated *Tg(dazl-ef1α:gfpd2)*^*y603*^ EpiTag transgenic animals at 5 dpf. Scale bar, 200 µm. (T) Quantification of the percentage of DMSO-treated and VPA-Treated *Tg(dazl-ef1α:gfpd2)*^*y603*^ EpiTag animals with bright GFPd2 livers.

We introduced the *dazl* CpG- or pCS vector sequence-containing transgenes into zebrafish embryos to generate EpiTag and control germline transgenic reporter zebrafish, respectively. As expected, the control transgenic line shows strong, ubiquitous expression of GFPd2 throughout early development (Fig. 1C-E and Fig. S2A-G). In contrast, EpiTag transgenic animals begin to express GFPd2 at around 6 hours post-fertilization, shortly after the MBT and zygotic gene activation. EpiTag GFPd2 fluorescence peaks at approximately 24 hpf and then gradually disappears thereafter, becoming essentially undetectable by 5 days post-fertilization (dpf) (Fig. 1F-H and Fig. S2H-N). GFPd2 silencing in the EpiTag line persisted into adulthood, where whole body (Fig. 2A,B) and organ (Fig. S3) GFPd2 expression is silenced whereas pCS control GFPd2 signal remains strong. The *Tg(dazl-ef1a:gfpd2)*^*y603*^ insertion was mapped to an intergenic region and would not be anticipated to affect gene function (Fig. S2O). We performed targeted bisulfite sequencing to examine whether *Xenopus ef1a* promoter methylation is influenced by neighboring *dazl* CGI or control vector sequences in the EpiTag and control pCS transgenic lines (Fig. 1I and Fig. S2P-S). In the pCS control line, methylation of the *Xenopus ef1a* promoter increased from approximately 7% to 11% between 1 and 5 dpf, while DNA methylation in the EpiTag line increased from 15% to 68% during the same time period (Fig. 1I and Fig. S2P-S). Increased DNA methylation of the *Xenopus ef1a* promoter in 5 dpf EpiTag larvae was accompanied by a decrease in activating H3K9 acetylation histone marks (Fig. 1J) and an increase in repressive H3K27me3 histone marks (Fig. 1K), as assessed by *Xenopus ef1a* chromatin immunoprecipitation (ChIP) assays. Together, these results suggest that the *dazl* CGI promotes methylation-induced epigenetic silencing of the adjacent *Xenopus ef1a* promoter.

**Figure 2.**
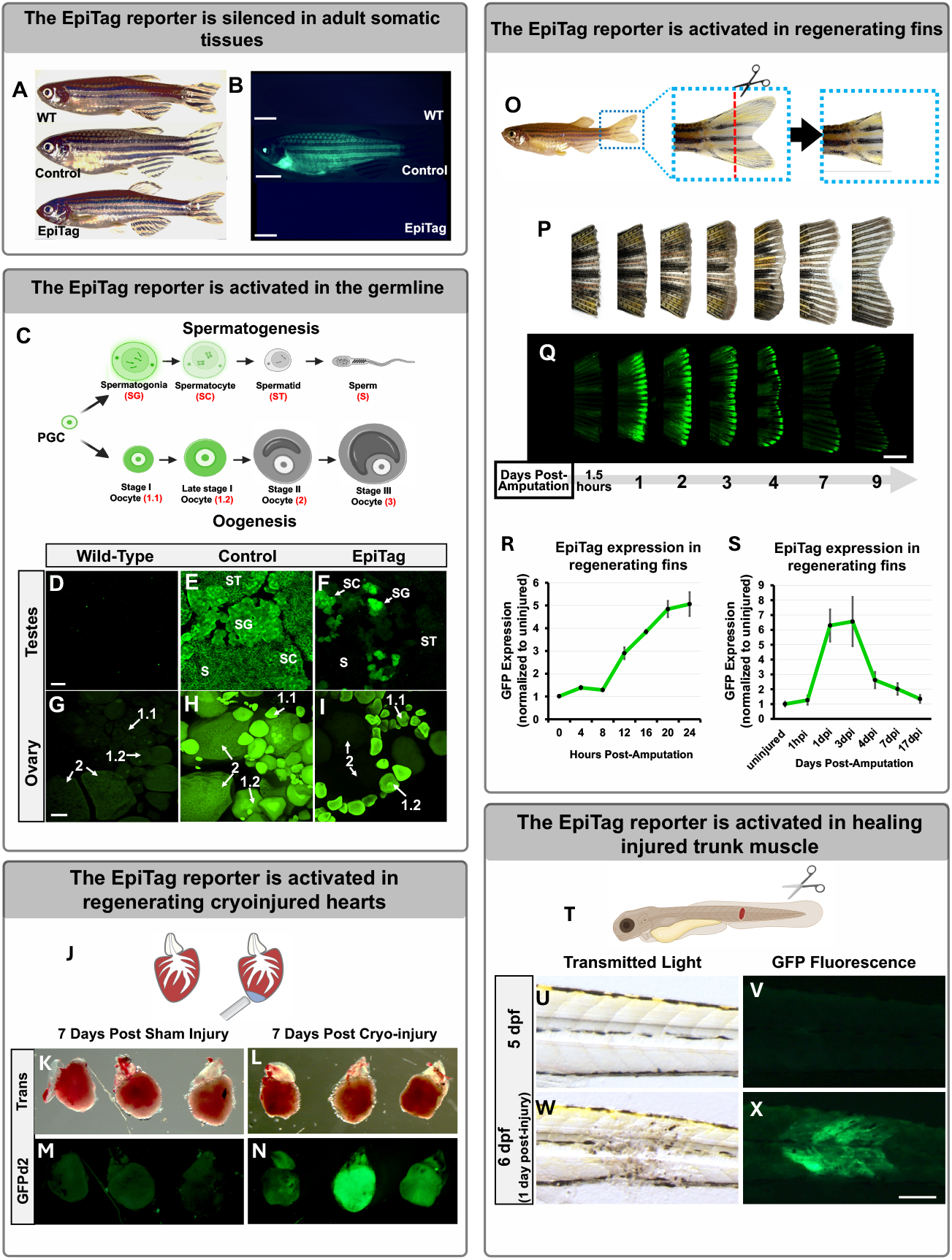
EpiTag reporter activation during gametogenesis and regeneration reprogramming. (A, B) Lateral view bright field (A) and green epifluorescence (B) images of intact wild-type (WT), *Tg(pcs-ef1α:gfpd2)*^*y605*^ Control, or *Tg(dazl-ef1α:gfpd2)*^*y603*^ EpiTag adult zebrafish. Scale bar, 0.5 cm. (C) Schematic diagram showing different stages of oogenesis and spermatogenesis starting from the primordial germ cell (PGC), with green coloring noting the corresponding GFPd2 expression patterns in EpiTag fish gonads. (D-F) Whole-mount ovary tissues of wild-type (D), *Tg(pcs-ef1α:gfpd2)*^*y605*^ Control (E), and *Tg(dazl-ef1α:gfpd2)*^*y603*^ EpiTag (F) females. Scale bar, 20 µm. (G-I) Whole-mount testes tissues of wild-type (G), *Tg(pcs-ef1α:gfpd2)*^*y605*^ Control (H), and *Tg(dazl-ef1a:gfpd2)*^*y603*^ EpiTag adult males. Scale bar, 20 µm. (J) Schematic of the cryoinjured heart procedure used for EpiTag expression analysis. Representative trans (K) and GFPd2 epifluorescence (M) images of dissected hearts seven days post-sham injury from adult EpiTag fish and trans (L) and GFPd2 epifluorescence (N) images of dissected hearts seven days post-cryoinjury from EpiTag fish. Scale bar, 100 µm. (O) Representative lateral view images of an EpiTag fish and its tail before and after clipping. (P,Q) Lateral bright field (P) and GFP confocal (Q) images of the regenerating tail of the same fin-clipped adult EpiTag zebrafish at 1.5 hours and 1, 2, 3, 4, 7, and 9 days post-amputation. Scale bar, 300 µm. (R,S) Quantitative RT-PCR analysis of *gfpd2* mRNA transcript levels normalized to the 0 hour or uninjured control levels every four hours for the first 24 hours post-amputation (R) and at 1 hour and 1, 3, 4, 7, and 17 days post-amputation (S). (T) Schematic diagram showing larval muscle injury protocol. (U-X) Representative bright field (U,W) and green epifluorescence (V,X) images of the trunk of the same EpiTag larva at 5 dpf just prior to trunk injury (U,V) and at 6 dpf, one day post-injury (W,X). Scale bar, 50 µm.

To assess the usefulness of the EpiTag line as an epigenetic reporter, we examined whether GFPd2 expression is affected by treatments known to disrupt epigenetic silencing. The DNA methyltransferase DNMT1 is required to maintain DNA methyl marks (9). Morpholino-mediated knockdown of zebrafish *dnmt1* in the EpiTag line leads to increased expression of GFPd2 at 2 dpf (Fig. 1L-N). 5-azacytidine (AZA) is a chemical analogue of the nucleotide cytosine that inhibits DNA methyltransferases and results in DNA hypomethylation (6). Treatment of EpiTag animals with AZA from the 256-512 cell stage enhances GFPd2 expression at 30 hpf compared to DMSO-treated control siblings (Fig. 1O-Q). Histone deacetylase 3 (Hdac3) is an epigenetic regulator thought to act as a co-repressor recruited to promoters regulated by nuclear hormone receptors and other transcription factors, where it represses transcription at least in part by removing “activating” histone modifications such as H3K9 acetylation. Zebrafish *hdac3* is expressed in the liver and is required for proper liver development (10). EpiTag animals treated from 1 dpf with the HDAC3-targeting class I HDAC inhibitor Valproic Acid (VPA) display liver-specific activation of GFPd2 expression at 5 dpf (Fig. 1R-T). Epigenetic changes can also be induced by certain environmental toxins, including nicotine, which causes aberrant DNA methylation (11, 12). Treating EpiTag larvae with nicotine results in GFPd2 activation at concentrations well below those resulting in lethality (Fig. S4). Together, these results suggest that the EpiTag line serves as an effective dynamic reporter of epigenetic silencing and activation.

Epigenetic reprogramming is known to take place during normal germline development (13) and the endogenous *dazl* gene is expressed during gametogenesis, so we examined whether GFPd2 is expressed in the gonads of adult EpiTag zebrafish and exhibits any stage-specific expression patterns in germline cells (Fig. 2C-I). Although the EpiTag reporter is silenced in somatic organs and tissues in wild-type (WT) adult fish (Fig. 2A, B), GFPd2 is actively expressed in both the male and female developing germline (Fig. 2C, F, I) at stages of spermatogenesis and oogenesis during which epigenetic reprogramming is known to take place (11). In contrast, WT ovaries and testes exhibit no GFPd2 expression (Fig. 2D, G) and pCS control ovaries and testes have strong GFPd2 expression in sperm (Fig. 2E) and oocytes (Fig. 2H) at all stages. Vasa is a well-known marker for early spermatogonia and oogonia cells, so we co-stained testes and ovary tissues with *vasa* and *gfpd2 in situ* hybridization chain reaction (HCR) probes to test their colocalization. Adult females exhibit EpiTag GFPd2 expression colocalized with *vasa* in stage 1 and 1.2 oocytes, but GFPd2 is undetectable in oocytes larger than approximately 150 µm in diameter (Fig. S5C), wheras adult EpiTag males exhibit GFPd2 expression in *vasa*-positive spermatagonia cells (Fig. S5H). Sectioned testes co-stained with antibodies recognizing Vasa, GFPd2, and Scyp3, another marker for spermatogonia and spermatocytes undergoing meiosis I and II, show that EpiTag GFPd2 and Scyp3 co-stain a subset of earlier spermatogonia and spermatocytes, but not later stage spermatocytes (Fig. S6C, S6F), unlike control pCS testes which exhibit GFPd2 at all stages (Fig. S6B, S6E). Thus, the EpiTag line serves as a useful marker of early stages of germ cell development in both testes and ovaries.

Epigenetic reprogramming is also known to take place during regeneration (14, 15). Zebrafish have a well-documented robust capacity to regenerate many different organs and tissues (16), and epigenetic reprogramming has been demonstrated in regenerating cells of many of these different organs and tissues (17). The EpiTag reporter is a very early and strongly activated marker of regenerating cells in several different commonly used zebrafish regeneration models (Fig. 2J-X and Figs. S7-S8). Zebrafish are able to regenerate their hearts following partial amputation or cryo-injury. Strong upregulation of the EpiTag reporter was observed in cryo-injured hearts at 7 days post-injury (Fig. 2J-N), with colocalization of GFPd2 and the regeneration marker PCNA noted in hearts with localized, limited cryoinjuries (Fig. S7). Caudal fin regeneration is probably the most widely used zebrafish regeneration model, with full regrowth of amputated fins occurring within a few weeks (Fig. 2O,P). The EpiTag reporter is strongly activated at the regenerating caudal fin margin by one day post-amputation (dpa), with expression declining thereafter essentially to background levels by 9 dpa (Fig. 2O-Q). EpiTag GFPd2 expression is particularly evident in and adjacent to the severed fin ray bones of 1 dpa fish, with strong expression in the emerging blastema and cellular-level resolution of GFPd2-positive cell types (Fig. S8A,B). As GFPd2 protein-folding and fluorescence is delayed by 30 minutes (18) to even 4 hours after the initiation of translation (19), we analyzed *gfpd2* transcript levels every 4 hours post-fin amputation (hpa) and detected increased *gfpd2* mRNA expression as early as 12 hpa in regenerating fin tissues (Fig 2R), with peak expression occurring at 3 dpa (Fig. 2S). Thus, we are able to use EpiTag expression to capture epigenetic reprogramming at stages much earlier than previously reported during zebrafish fin regeneration (20, 21). Zebrafish also actively repair and regenerate injured skeletal muscle (22). Injury of skeletal muscle in the trunk of larval EpiTag zebrafish (Fig. 2T-X) results in strong activation of the GFPd2 in cells at and closely adjacent to the wound site within a day after injury (Fig. 2W, X, Fig. S8H, J) compared with wild-type injured embryos which only exhibit weak green pigment cell autofluorescence in the trunk (Fig. S8G, I). These results indicate that the EpiTag reporter is strongly activated in and marks cells participating in regenerative processes. Together, our findings suggest the EpiTag reporter is a useful tool for directly visualizing cells, tissues, and organs undergoing gametogenesis or regenerative processes in living zebrafish.

Forward-genetic mutagenesis screens in invertebrates have been highly successful in identifying key players in epigenetic regulation in these animals (23-25). We reasoned that the EpiTag zebrafish might provide a useful tool for a similar forward-genetic analysis to uncover vertebrate global or tissue-specific epigenetic regulators of developmental programs. We used the EpiTag line to carry out an F3 ENU mutagenesis screen for recessive mutants exhibiting activation or suppression of GFPd2 reporter expression (Fig. 3A), screening for mutants either prematurely silencing GFPd2 at 2 dpf or displaying ectopic ubiquitous or tissue-specific expression of GFPd2 at 5 dpf. We screened 110 F2 families (220 mutagenized genomes) derived from 14 mutagenized founders and identified 7 mutants with either decreased GFPd2 expression at 2 dpf or increased GFPd2 expression at 5 dpf (Table S1). Two mutants show reduced expression at 2 dpf (Table S1, fig. s9E, H), with two of these five mutants also showing increased trunk GFPd2 expression at 5 dpf (Table S1). Five mutants show organ- or tissue-specific activation of GFPd2 expression at 5 dpf (Table S1), with different mutants showing expression in the liver (Fig. S9J, K), pharynx, blood, and brain (Table S1). RNAseq-based bulked segregant mapping (26, 27); see methods) was used to map EpiTag mutants to narrowly defined intervals on specific chromosomes of the zebrafish genome (Fig. S9A). The RNAseq sequencing data was then used to narrow the list of genes within each mapped interval to the most likely candidate(s) for the defective gene by (i) identifying coding sequences with potentially deleterious lesions (i.e., termination, defective splicing, or amino acid changes), (ii) uncovering evidence of gene expression in the relevant organ or tissue, and/or (iii) gene annotations suggesting potential function as or interaction with known epigenetic regulators. We conducted further in-depth characterization of a “liver GFP on” y595 mutant. This mutant shows very strong, specific expression of GFPd2 in the liver at 5 dpf (Fig. 3B-E), suggesting the defective gene is influencing epigenetic programs in an organ-specific manner. The mutant maps to an interval on chromosome 20 containing *phosphoribosyl pyrophosphate amidotransferase* (*ppat*; Fig. 3F,G), the zebrafish ortholog of a gene encoding an enzyme which catalyzes the first step in the *de novo* purine nucleotide biosynthesis pathway (Fig. 3J). In y595 mutants, *ppat* contains a G to T mutation in exon 10, resulting in a nonsynonymous C426F amino acid change (Fig. 3G, H) in a putatively important enzymatic binding site. Whole mount *in situ* hybridization of 3 dpf zebrafish shows enriched specific expression of the *ppat* gene in the developing liver (Fig. 3I), with continued liver-specific expression in 4-5 dpf fish (Fig. S10). To confirm *ppat* is responsible for the y595 mutant phenotype, we used CRISPR/Cas9 technology to create a second targeted allele of *ppat* (*ppat*^*y710*^) with a 4 base deletion spanning nucleotides 25,637,783 – 25,637,786 in exon 10, only 10 to 13 nucleotides upstream of the y595 SNP allele, resulting in a frameshift and predicted truncated polypeptide (Fig. 3H, Fig. S11). The *ppat*^*y710*^ mutant displays a similar GFPd2 activation in the liver and fails to genetically complement the original *ppat*^*y595*^ mutant (Fig. S11), providing strong evidence the *ppat* gene lesion is in fact the causative defect for the *ppat*^*y595*^ phenotype. Due to the importance of Ppat in *de novo* purine synthesis (Fig. 3J), we investigated whether *ppat*^*y595*^ mutants had altered levels of two downstream metabolites of the pathway, guanine and uric acid. Biochemical assays confirmed that *ppat*^*y595*^ mutants had drastically reduced levels of both uric acid (Fig. 3K) and guanine (Fig. 3L), confirming *ppat*^*y595*^ is a hypomorphic allele of *ppat*. To determine whether uric acid or guanine deficiencies were responsible for the liver GFPd2 phenotype, we selectively abolished *xdh* or *pnp* (Fig. 3J) gene expression via CRISPR/Cas9 dgRNP-mediated techniques (Fig. S12). Targeting guanine via *pnp* dgRNP injections (Fig. 3N), but not uric acid via *xdh* dgRNP injections (Fig. 3M), resulted in liver GFPd2-positive embryos that phenocopied *ppat* mutants (Fig. S12). These findings suggest the liver-specific epigenetic activation is a result of purine nucleotide, but not uric acid, imbalance. Interestingly, *ppat*^*y595*^ and *ppat*^*y710*^ mutants also exhibited a lack of xanthophore pigmentation at embryonic stages (Fig. S13A,B) despite having normal development and expression of xanthophore cell markers (Fig. S13I-N). Homozygous *ppat*^*y595*^ mutants survived to adulthood, and did not exhibit xanthophore pigmentation defects at this stage (Fig. S13C-H), although they did exhibit major fin abnormalities (Fig. S14).

**Fig. 3.**
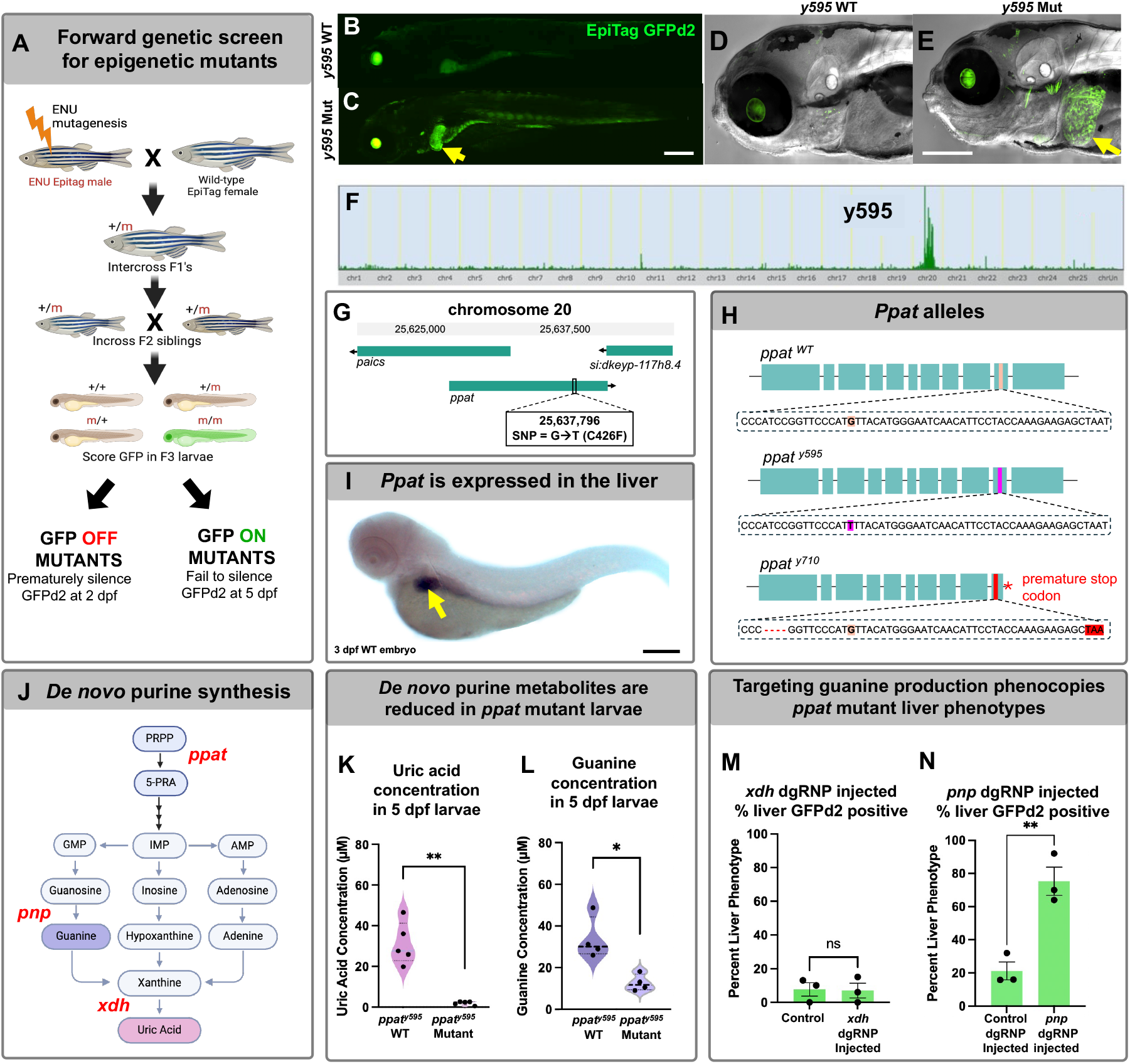
Isolation of tissue-specific epigenetic mutants using EpiTag transgenic animals reveals a liver-specific EpiTag activation mutant with a defect in *ppat*. (A) Schematic diagram showing the design of an F3 ENU mutagenesis screen using EpiTag transgenic fish to isolate recessive mutations that either prematurely silence GFPd2 expression at 2 dpf or fail to silence GFPd2 expression at 5 dpf. (B,C) Representative GFP confocal images of y595 wild-type phenotypic larvae (B) and phenotypic mutant siblings (C) exhibiting liver-specific GFPd2 expression at 5 dpf. Scale bar, 200 µm. (D,E) Higher magnification DIC and GFP confocal images of y595 wild-type (D) and mutant (E) larvae exhibiting bright GFPd2 expression in liver tissues (arrow). Scale bar, 200 µm. Green eyes represent autofluorescence. (F) Tracks for the Hidden Markov Model (HMM) SNPTrack scores identifying regions of interest of the y595 mutant genome SNPs compared with WT siblings. (G) Representative image of the chromosome region where the gene *ppat* was found to exhibit a SNP resulting in a cytosine (C) to phenylalanine (F) missense mutation in mutant larvae. (H) Schematic depicting the *ppat* exons and affected regions in WT, y595, and y710 alleles. (I) Standard *in situ* hybridization of a 3 dpf WT embryo where *ppat* shows strong specific expression in liver tissues (arrow). Scale bar, 200 µm. (J) Schematic diagram of the *de novo* purine synthesis pathway of which Ppat (red text) catalyzes the first enzymatic reaction. (K, L) Quantification of guanine (K) and uric acid (L) concentrations present in pooled 5 dpf *ppat*^*y595*^ WT and mutant siblings. (M, N) Quantification of the percentage of 5 dpf wild-type EpiTag fish exhibiting GFPd2 positive livers after *xdh* dgRNP injections (M) or *pnp* dgRNP injections (N) compared with controls.

Due to the liver-specific expression of *ppat* and epigenetic activation in these tissues, we examined liver phenotypes in adult *ppat* mutant and WT sibling fish and discovered that mutant livers exhibited strongly increased intracellular lipid droplet accumulation as seen via histological analyses (Fig. 4A, B) and lipid-specific stains (Fig. 4C-E). Adult *ppat* mutant livers also exhibited other fatty liver disease symptoms, including ballooned hepatocytes (Fig. S15), indicative of unhealthy liver cells. To determine whether *ppat*^*y595*^ mutants exhibited liver-specific transcriptomic or methylomic signatures related to their liver disease pathology, we conducted RNASeq and whole genome bisulfite sequencing (WGBS) on the same adult liver tissues from genotyped y595 homozygous mutant, heterozygote, and wild-type sibling animals (Fig. 4F). Adult *ppat*^*y595*^ mutants exhibited distinct upregulation of many genes compared with heterozygote and WT counterparts (Fig. 4G), and the GO-terms associated with these genes related to many metabolic dysfunction-associated fatty liver disease phenotypes (MAFLD) (Fig. 4H). Many of the upregulated genes corresponded to human genes dysregulated in patients with MAFLD or advanced stages of MAFLD resulting in cirrhosis and hepatocellular carcinoma (Fig. 4I). Transcriptomic/methylomic profiling was also performed on larval liver tissues to look for early changes that might predispose *ppat* mutants to MAFLD (Fig. 4J). RNASeq and WGBS of FACS-sorted 6 dpf *ppat* mutant and phenotypically WT sibling larval liver cells revealed upregulation of several genes related to liver lipid metabolism and pathologies (Fig. 4K). We further validated the liver-specific upregulation (Fig. 4L,N) of two of the lipid metabolism genes, *slc7a3a* and *adm2a*, by using whole-mount HCR (Fig. 4P-S) or standard *in situ* hybridization (Fig. S16) to demonstrate liver-specific upregulation of their transcripts. Increased expression of *slc7a3a* and *adm2a* persisted in adult liver tissues (Fig. 4M, O), supporting the idea that early molecular changes occurring in *ppat* mutant livers may be contributing to the progression of liver disease later in life.

**Figure 4.**
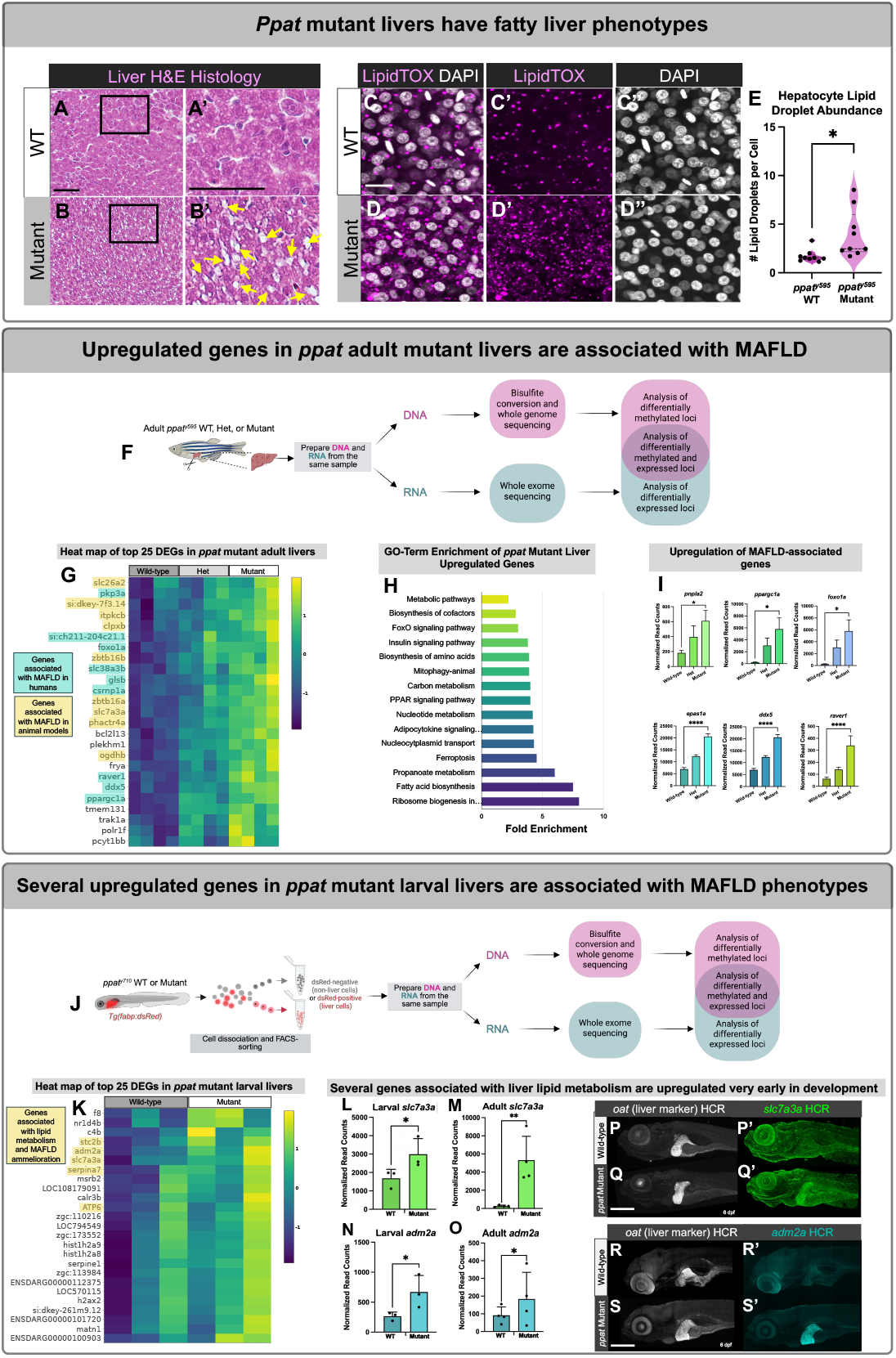
*Ppat* mutant zebrafish livers exhibit metabolic fatty liver disease physiological and molecular phenotypes. (A,B) Histological hematoxylin and eosin (H&E) stains of wild-type (A) and *ppat*^*y595*^ mutant adult liver sections with lipid droplets highlighted (yellow arrows). Scale bar, 50 µm. (C, D) Liver tissues stained with LipidTox dye from adult *ppat*^*y595*^ WT (C) and mutant (D) sibling fish. Scale bar, 25 µm. (E) Quantification of the average number of lipid droplets present per cell from *ppat*^*y595*^ WT and mutant fish. (F) Schematic diagram of the experimental pipeline for tissue isolation and RNASeq and whole genome bisulfite sequencing (WGBS) of the same adult liver tissues from adult *ppat* homozygous mutant, heterozygote, and wild-type (WT) sibling liver tissues. Samples were prepared and sequenced in triplicate. (G) Heat map showing the top 25 upregulated genes in *ppat* mutant livers and their fold change expression differences compared with heterozygote and WT livers, with genes associated with MAFLD in humans highlighted in blue and MAFLD in other animal models highlighted in yellow. (H) Gene ontology enrichment of loci significantly upregulated in *ppat* mutant livers compared with heterozygote and WT counterparts. (I) Graphs depicting representative genes which have been associated with human patients with metabolic dysfunction-associated fatty liver disease (MAFLD) and their RNASeq normalized read count fold-change differences in *ppat* mutant, heterozygote, and WT liver tissues. (J) Schematic diagram of the experimental pipeline to isolate *ppat* mutant and WT larval liver tissues and subsequent RNASeq and WGBS analysis. Samples were prepared and sequenced in triplicate. (K) Heat map of top 25 differentially expressed genes upregulated in *ppat* mutant larval liver tissues with the genes associated with lipid metabolism and/or MAFLD highlighted in yellow. (L, M) Normalized read counts of *slc7a3a* which shows upregulation in larval (L) and adult (M) liver tissues along with *adm2a* which shows upregulation in larval (N) and adult (O) liver tissues. (P, Q) *In situ* HCR of *slc7a3a* and a control liver gene (*oat*) in 6 dpf ppat WT (P) and mutant (Q) larvae. Scale bar, 200 µm. (R, S) *In situ* HCR of *adm2a* and a control liver gene (*oat*) in 6 dpf ppat WT (R) and mutant (S) larvae. Scale bar, 200 µm.

We also conducted WGBS analysis on both adult and larval liver tissues to determine if any changes in DNA methylation were occurring in liver tissues due to the liver-specific epigenetic activation in *ppat* mutants, and if any differentially methylated loci were associated with MAFLD-associated gene expression changes. In adult liver tissues many MAFLD-associated genes also exhibited hypomethylation in mutants compared to wild-type or heterozygote livers, suggesting differences in methylation may be contributing to the expression differences in these genes (Fig. 5A,B, Fig. S17). We also discovered two histone demethylases previously implicated in hepatocyte lipid metabolism (28), Kdm2aa and Kdm6ba, exhibited both increased expression in *ppat* mutant livers and differential methylation patterns (Fig. S18), suggesting that purine nucleotide imbalance leads to differences in liver-disease associated histone methylation as well.

**Figure 5.**
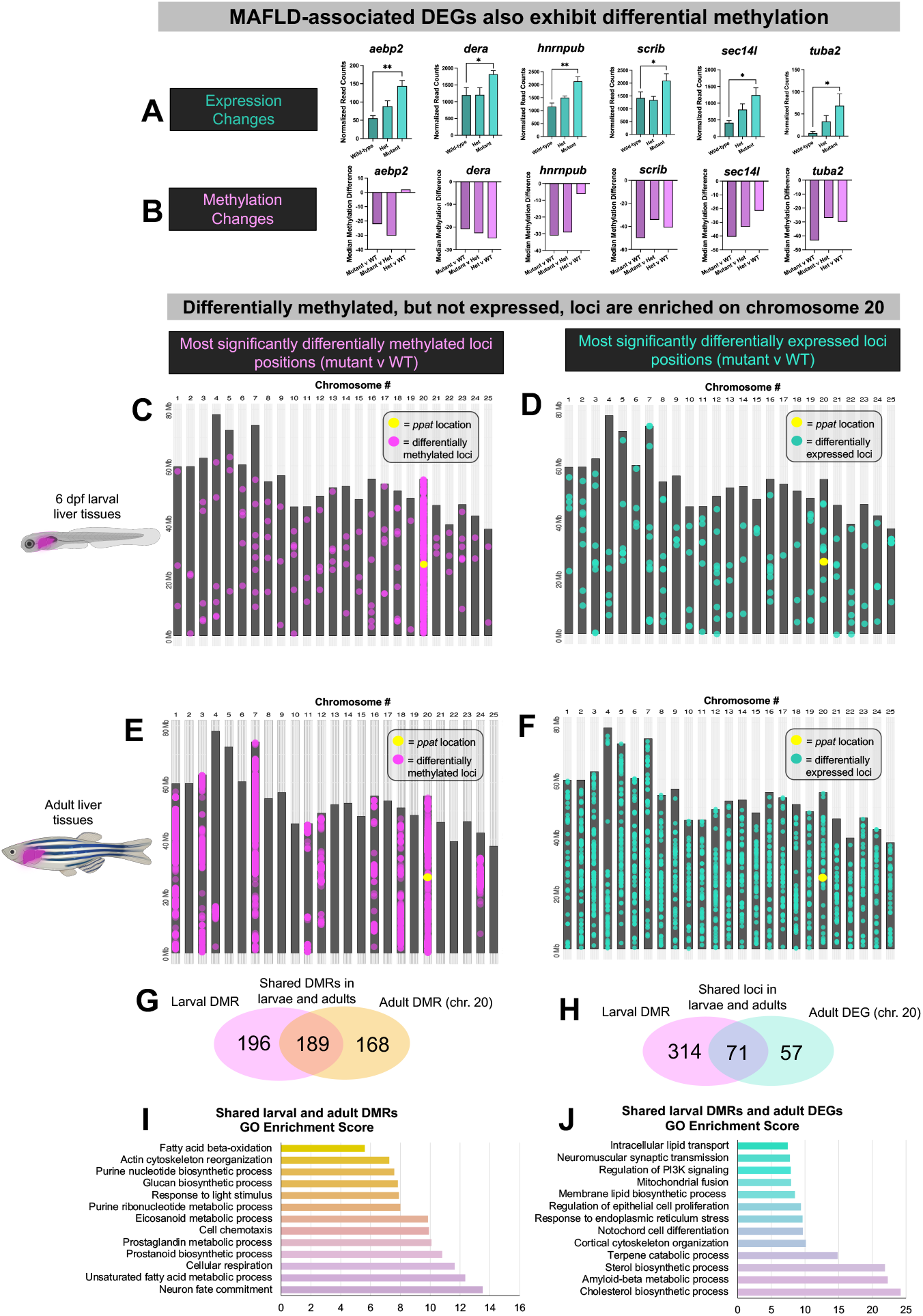
*Ppat* mutant differentially methylated loci are significantly enriched in chromosome 20, overlap with later differentially expressed genes, and are associated with fatty acid metabolic functions. (A,B) Representative genes with relevance to metabolic dysfunction-associated fatty liver disease (MAFLD) which show upregulation in *ppat* adult mutant livers (A) and hypomethylation in comparisons of mutant and WT or mutant and heterozygote tissues (B). (C,D) Chromosomal maps depicting the locations of the most significant differentially methylated loci (q-value < 0.05, magenta dots) in *ppat* mutant larvae compared with WT (C), and the most significantly differentially expressed genes (p-adj < 0.05, teal dots, D). (E, F) Chromosomal maps depicting the locations of the 10,000 most significant differentially methylated loci (q-value < 1.485E-15, magenta dots) in *ppat* mutant adult livers compared with WT (E), and the most significantly differentially expressed genes (p-adj < 0.05, teal dots, F). (G) Venn diagram depicting the unique and shared number of significantly differentially methylated loci in larval and adult livers. (I) Gene set enrichment analysis of shared loci depicted in G. (H) Venn diagram displaying the unique and shared number of significantly differentially methylated loci in *ppat* mutant larvae and differentially expressed genes (DEGs) in *ppat* mutant adult livers. (J) Gene set enrichment analysis of shared loci depicted in H.

Remarkably, the majority of the differentially methylated loci identified in larval *ppat* mutant livers were located on chromosome 20 (2,514 loci out of 2,618 loci) (Fig. 5C), the same chromosome where *ppat* is located, whereas differentially expressed genes were scattered across all chromosomes (Fig. 5D). This chromosome 20 enrichment of methylated loci persisted in adults, where chromosome 20 was the second-most enriched for differentially methylated loci, with chromosome 7 and several others also exhibiting strong enrichment in differentially methylated loci (Fig. 5E). Adult significantly differentially expressed genes again did not display chromosomally enriched patterns, unlike the methylated loci (Fig. 5F). *Ppat* mutant adult and larval livers shared 189 differentially methylated loci (Fig. 5G), which was highly statistically above the expected shared number of loci (Fig. S19A). GO terms associated with these loci were related to lipid metabolic pathways and other pathways with relevance to MAFLD (Fig. 5I). We also investigated whether early epigenetic changes were associated with later expression changes in liver tissues, comparing the shared loci that were both differentially methylated in larvae and differentially expressed in adult livers. There was a much higher number of these shared loci than would be expected statistically (Fig. 5H, Fig. S19B), again with loci-associated GO terms involved in lipid metabolism, indicating that early differences in methylation at these loci may be contributing to the later fatty liver phenotype. We investigated a few of the specific genes, such as *med23* and *cyp2v1*, that showed differential larval promoter methylation associated with increased (Fig. S20A, B) or decreased (Fig. S20D, E) expression in adults. *Med23* encodes for an epigenetic transcriptional regulator which promotes liver adipogenesis and glucose metabolism via upregulation of *Foxo1* expression (29), whereas *cyp2v1* is an ortholog of mammalian *Cyp2j2*, which has been shown to attenuate MAFLD disease via its hepatoprotective roles (30). The differential methylation and expression of these genes may in part contribute to the progression of MAFLD phenotypes in *ppat* mutant fish.

Although PPAT has not been previously implicated specifically in epigenetic regulation, our findings indicate that variants in this gene lead to profound dysregulation of methylation, tissue-specific defects in epigenetic silencing, and development of fatty liver disease in zebrafish. Future studies further exploring potential crosstalk between purine biosynthesis variants and epigenetic regulation may lead to new insights into links between metabolism, epigenetics, and disease pathologies. Together, our findings demonstrate the power of EpiTag zebrafish for *in vivo* visualization, experimental analysis, and genetic dissection of epigenetics and its role in vertebrate development, gametogenesis, regeneration, and disease.

## Supporting information

Supplemental Data Part 1

Supplemental Data Part 2

## Acknowledgments

We thank Dr. Katie Drerup for help with ENU mutagenesis and Drs. Karuna Sampath, Ajay Chitnis, Igor Dawid, Jeff Gross, Mary Goll, Elke Ober and the zebrafish community for sharing reagents, protocols and discussions. We also thank Kelly Tomins, Iana Sahadzic, Jeremy Otridge, Charlotte Visconti, Catie Bohmer and Alisa Forsberg for their help with the ENU mutagenesis screen. We thank Drs. Leonard Zon and Marlies Rossmann for sharing unpublished observations. We apologize to those whose original work could not be cited because of space limitations. Funding: This research was supported by the intramural programs of the NICHD grant ZIA-HD008977 (to BMW). Author contributions: A.V.G., K.T., M.M., and D.C. participated in the ENU mutagenesis screen. M.M., A.V.G., K.T., D.C., G.M., A.G, A.A.S, O.C., J.C., M.D., L.G., A.P., Y.A.K., J.M.K., A.E.D, M.K., J.P., K.B., J.I., T.L., F.F., R.D., and B.M.W. designed and performed the experiments and analyzed the data. M.M, A.V.G., and B.M.W. wrote the manuscript with inputs from all the authors. Competing interests: The authors declare no competing interests. Data and materials availability: All data needed to reproduce and evaluate the conclusions in this paper are present in the main text and supplementary materials.

## Materials and Methods

### Zebrafish husbandry and strains

Fish were housed in a large zebrafish dedicated recirculating aquaculture facility (4 separate 22,000 L systems) in 6L and 1.8L tanks. Fry were fed rotifers and adults were fed Gemma Micro 300 (Skretting) once per day. Water quality parameters were routinely measured, and appropriate measures were taken to maintain water quality stability (water quality data available upon request). Zebrafish lines used in this study are maintained and staged as described previously (1, 2). Zebrafish lines used in this study are *Tg(dazl-ef1a:gfpd2)*^*y603*^, *Tg(pcs-ef1a:gfpd2)*^*y605*^, *ppat*^*y595*^, *ppat*^*y710*^ (CRISPR mutant; 4 bp deletion), *Tg(fabp:dsRed)* (3), and EK and TL as wild-type. This study was performed in an AAALAC accredited facility under an active research project overseen by the NICHD ACUC, Animal Study Proposal # 24-015.

### Cloning of the transgenic constructs and generation of stable transgenic lines

Constructs were assembled using vectors from the Gateway Tol2kit (4). In brief, the 5’ promoter CpG island from the *dazl* gene and pCS vector sequences were cloned separately into the 5’ entry vector. The CpG island located in the 5’ region of the *dazl* locus on Chr. 19 from 2,04,02,938 to 2,04,03,755 was amplified using the PCR primers (see Table S2 for primer details). The *Xenopus ef1a* promoter and *gfpd2* were cloned into middle entry and 3’ entry clones respectively. A final construct was assembled using pDestTol2pA2 from the Tol2kit (4). All constructs were sequenced prior to injections. For generation of stable transgenic lines, 1-cell stage fertilized embryos derived from natural matings of wild-type (EK strain) zebrafish were injected with either *Tg(dazl-ef1a:gfpd2)*^*y603*^ or *Tg(pcs-ef1a:gfpd2)*^*y605*^ constructs along with transposase RNA. Embryos were screened for GFPd2 expression at 1 and 2 dpf. GFPd2 positive embryos were raised to adulthood and crossed to screen for germline carriers. GFPd2 positive F1 embryos were raised to adulthood and used to generate stable transgenic lines on the WT EK background. We mapped the insertion sites as previously described (5).

### Chemical mutagenesis and genetic screen

Homozygous *Tg(dazl-ef1a:gfpd2)*^*y603/y603*^ “EpiTag” males on the EK background were used for N-ethyl-N-nitrosourea (ENU) mutagenesis. ENU mutagenesis was carried out as described previously with minor modifications (6). Four rounds of ENU mutagenesis were performed at one-week intervals to allow mutagenized fish to recover. F1 families were generated by IVF using mutagenized males and homozygous EpiTag females on the EK background. F2 families were generated by in-crossing F1 fish. F3 larvae derived from crossing pairwise F2 family fish were screened for premature loss or persistent presence of GFPd2 expression at 48 hpf and 5 dpf, respectively. We screened at least 50% of the F2 fish per family (typically 8-14 independent pairs per F2 family). Pairs which showed 25% of the embryos from the clutch with the presence or absence of GFPd2 expression were isolated as mutant carriers. For mapping crosses, mutant carriers were crossed to an EpiTag-TL mapping line. The EpiTag-TL mapping line was generated by backcrossing EpiTag-EK background fish to wild type TL strain fish for six generations. Mapping pairs were identified from these hybrid background fish and used subsequently for mapping the mutant gene. For mapping, mutant and sibling embryos (at 2 or 5 dpf, based on when the mutant phenotype was seen) derived from the same pair of parent fish were collected after screening for the phenotype. Two separate pools of 20-50 mutant and 20-50 phenotypically wild type sibling embryos were used for total RNA extraction. Extracted RNA was used for RNAseq. Typically, we used 30-40 million PE reads for mapping. Fastq files were used in RNAmapper (https://genome.cshlp.org/content/23/4/679), SNPTrack (http://genetics.bwh.harvard.edu/snptrack/), or a modified version of RNAmapper. We tested each RNAseq dataset with at least two mapping programs to verify the predicted linked chromosomal interval. The RNAseq datasets included a variety of SNPs in the mapped intervals that were enriched in the mutant RNAseq data set. These SNPs were further filtered based on whether the SNP was gene-associated, the type of gene mutation (stop codon, nonsynonymous mutation, synonymous mutation), the expression pattern of the gene, and previous evidence that the gene interacts with epigenetic regulators. If required, we performed additional fine mapping using the SNP information derived from the RNAseq mapping data to narrow down the linked region and identify the mutated gene.

### Targeted mutation generation using CRISPR/Cas9

Mutations in the zebrafish *ppat* gene was generated using the CRISPR/Cas9 system. gRNA targets and screening primer pairs were selected from GRCz11 assembly using CHOPCHOP (7). Cas9 and gRNAs were injected in the fertilized one cell stage EpiTag (*Tg(dazl-ef1a:gfpd2)*^*y603*^) embryos. To test the efficiency of RNA targeting, embryos were collected at 1 dpf and assayed for indel formation using fluorescent PCR and fragment analysis on an ABI 3730 DNA Analyzer. The *ppat* locus on chromosome 20 was targeted at exon 10 using the guide RNAs sequences listed in Table S2. To establish stable mutant-carrying lines, embryos were injected with Cas9 and gRNAs and then raised to adulthood. To identify fish transmitting germline mutations, the injected adults were crossed *to Tg(dazl-ef1a:gfpd2)*^*y603*^ fish and the embryos were collected at 1 dpf and assayed for segregation of specific indels using fluorescent PCR and fragment analysis on an ABI 3730 DNA Analyzer. Additional embryos were raised to adulthood to establish germline mutant lines. We identified an allele of *ppat* with a 4 bp deletion in exon 10 (*ppat*^*y710*^); this allele is predicted to induce a premature stop codon and early truncation of the Ppat peptide in the same region as the SNP mutation.

### Morpholino injection and drug treatments

The *dnmt1* morpholino 5′-ACAATGAGGTCTTGGTAGGCATTTC-3′ was purchased from Gene Tools. Either 1 or 2 ng of *dnmt1* morpholino was injected into one cell stage fertilized embryos. 5-Azacytidine (Aza) was purchased from Sigma and dissolved in DMSO to prepare a 10 mM stock solution. EpiTag embryos were treated in 50 µM Aza in embryo media from blastula stage (2-2.5 hpf) until 24-30 hpf and then assayed for phenotypes. Valproic acid (VPA) was purchased from Enzo and dissolved in DMSO to prepare a 10 mM stock solution. EpiTag embryos were treated with 5 µM VPA from 1-5 dpf and then assayed for phenotypes. Nicotine was purchased from Sigma and diluted in embryo medium to final concentrations from 10 µg/ml to 200 µg/ml.

### DNA/RNA extraction and RNA sequencing for SNP mapping

For mapping SNP mutants, DNA and RNA were co-isolated using a Zymo Quick DNA/RNA Microprep Plus Kit (Zymo Research; D7005). For RNA sequencing, poly-A enriched RNA was converted to indexed libraries from each sample (sibling and mutant pools) using the TruSeq mRNA library prep Kit (Illumina). Libraries were sequenced on a HiSeq 2500 to generate >100 million paired end reads (100 bp) per sample. Resulting Fastq files were used for mapping the mutants.

### RNA/DNA isolation, library preparation, and sequencing for dual RNASeq and WGBS

For dual RNASeq sequencing and whole genome bisulfite sequencing (WGBS), DNA and RNA were co-isolated from either whole dissected livers or FACS-sorted cells from 6 dpf larvae using the Quick DNA/RNA Microprep Plus Kit (Zymo Research; D7005). For RNASeq libraries, total RNA was reverse transcribed to cDNA, cDNA from rRNA depleted, and dual indexing conducted using Zymo-Seq™ RiboFree Total RNA Library Kit (Zymo Research; R3000). For WGBS libraries, bisulfite conversion, amplification, and indexing was conducted using the Zymo-Seq™ WGBS Library Kit (D5465-A) and tagmentation was conducted using the Illumina DNA Prep, (M) Tagmentation kit (20060060). RNASeq and WGBS libraries were quantified and quality checked with an Agilent 2100 Bioanalyzer using 1-2 nL of each library sample with the RNA 600 Pico Kit (cat no. 5067-1513, Agilent) or the High Sensitivity DNA Assay Kit (Agilent; 5067-4626). RNASeq sequencing was conducted on a NovaSeq 6000 using a NovaSeq SP Reagent Kit v1.5 (200 cycles) flow cell to generate a minimum of 400 million 2 × 100 bp reads and WGBS sequencing was conducted on a NovaSeq 6000 using an S4 reagent kit v1.5 (200 cycles) to generate a minimum of 8 billion 2 × 100 bp reads.

### RNASeq Analysis

We used lcdb-wf pipeline (https://github.com/lcdb/lcdb-wf, version v1.10.3rc) to determine differentially expressed genes. The pipeline aligns the sequencing fastq files, performs gene expression quantification, differential gene expression and functional enrichment analyses (The main tools we used in the pipeline included hisat2 v2.2.1, subread v2.0.3, DESeq2 v1.38.0 and clusterProfiler v4.6.0.) The pipeline outputs html summary files and its output can be fed directly to an interactive app Carnation (https://github.com/NICHD-BSPC/carnation) which was used for data exploration. Gene set enrichment analysis was conducted using FishEnrichr (8, 9).

### Whole genome bisulfite sequencing (WGBS) Analysis

For WGBS analysis, raw paired-end fastq files were trimmed with trim-galore (v 0.6.10) with flags “--clip_r1 15 --clip_r2 15 --three_prime_clip_r1 1 --three_prime_clip_r2 1”. We then used Bismark suite of tools (v0.24.2) to align the trimmed fastq files. The following flags were used for alignment: “--bowtie2 -D 20 -R 3 --score_min L,0,-0.6 --non_bs_mm --nucleotide_coverage -- non_directional –dovetail”. Poorly bisulphite-converted fragments were subsequently filtered out with “filter_non_conversion -p --percentage_cutoff 20 --minimum_count 3” and the remaining fragments were deduplicated with deduplicate_bismark. Bismark_methylation_extractor was used to extract per-CpG methylation data for downstream analysis. We used methylKit (v1.28.0) to first determine differential methylation at the single-CpG level and then determine regions of differential methylation, based on segmentation. We retained segments with |mean methylation difference| > 10, together with their statistics and associated gene annotations. For larvae WGBS, we pooled all replicates of each condition due to insufficient and heterogeneous concerns in individual samples. Functional enrichment of gene promoter-associated regions was performed with clusterProfiler (v 4.10.0). Differentially methylated loci for larvae were selected for additional analysis, such as chromosomal mapping analysis, using differentially methylated loci regions with a best.q value < 0.05 and differentially methylated loci for adults were selected using the top 10,000 most differentially methylated which corresponded to loci with a best.q value < 1.485E-15.

### RT-qPCR and CUT&RUN

DNA and RNA were co-isolated using a Zymo Quick DNA/RNA Microprep Plus Kit (Zymo Research; D7005). Equal amounts of RNA were converted into cDNA using a ThermoScript RT– PCR System (Invitrogen). The resulting cDNA was used in quantitative PCR using SsoAdvanced Universal SYBR Green Supermix (Bio-Rad) on a CFX96 Real-Time system. A minimum of three biological replicates were performed. CUT&RUN was performed as described in the manufacturer’s instructions using protein A fused to micrococcal nuclease (pA-MNase) purchased from EpiCypher (15-1016). In brief, 1- and 5-dpf EpiTag and PCS-control embryos were dissociated to prepare single cell suspension as described above. Roughly 100-120 embryos were used in each assay. CHIP grade antibodies for H3K27 trimethylation (C36B11), H3K9Ac (C5B11) and total H3 (D1H2) were purchased from Cell Signaling Technology and used in CUT&RUN. Sequences of primers used for RT-qPCR and CUT&RUN are listed in Table S2.

### Imaging, whole mount in situ hybridization, whole mount in situ hybridization chain reaction, immunohistochemistry and AFOG staining

Live embryo imaging, whole mount *in situ* hybridization, and immunohistochemistry were performed as described previously (10, 11). Whole mount *in situ* hybridization chain reaction (HCR) on testes, ovaries, and larvae were performed as described previously (12). All HCR samples were stored in 5X saline sodium citrate solution with 0.1% Tween-20 (SSCT) in the dark and then imaged on a Nikon Ti2 inverted microscope with Yokogawa CSU-W1 spinning disk confocal (Hamamatsu Orca Fusion-BT camera). HCR probe sets for *vasa, gfpd2*, and *slc7a3a* were designed by Molecular Instruments (Molecular Instruments, Los Angeles, CA) and probe sets for *oat* and *adm2a* were designed in-house and ordered as 50 pmole oligo pools (oPool) from Integrated DNA Technologies. For immunohistochemistry, we cryo-sectioned zebrafish ovaries, testes, and hearts. Testes sections were stained using Chicken anti-GFP (Abcam; ab13970), Rabbit anti-Vasa (GeneTex; GTX128306), Rabbit anti-Sycp3 (Abcam; ab97672) primary antibodies. Ovary sections were stained using Chicken anti-GFP (Abcam; ab13970). For heart sections, we used Chicken anti-GFP (Abcam; ab13970) and Rabbit anti-PCNA (GeneTex; GTX124496) as primary antibodies. All sections were stained using anti-Chicken Alexa 488, anti-Rabbit Alex 546 secondary antibodies (ThermoFisher) and ToPro3 (ThermoFisher) as nuclear dye. We performed AFOG staining as described previously (13).

### Zebrafish injury assays

Adult zebrafish heart cryoinjury and fin amputations were performed as described previously (13, 14). In brief, for heart cryoinjury adult EpiTag zebrafish were anesthetized using 1X tricaine solution. A small incision was made in the ventral body wall to expose the heart. A probe cooled in liquid nitrogen was placed on the exposed ventricle until localized freezing was seen (a few seconds). Zebrafish were immediately transferred to a tank of fresh water to recover. Sham treatments were done similarly except that a room temperature probe was used. For tail fin injury, 40-50% of the tail fin of an adult anesthetized fish (EpiTag) was amputated using razor blade or iridectomy scissors. Fish were returned to a tank with fresh water. The same fish was imaged every day to monitor GFPd2 expression in the regenerating tail fin. Larval skeletal muscle injury was performed as described previously (15). Briefly, EpiTag and WT larvae were anesthetized in 1X tricaine then mounted in low melting point agarose and a small wound was made to the side of the mid-trunk using fine iridectomy scissors (1-3 somites). Larvae were removed from the agarose and allowed to recover in fresh embryo culture media, then analyzed 1 day post-injury for the presence of GFPd2 fluorescence.

### Larvae cell dissociation and FACS-sorting

Zebrafish larvae at the 6 dpf stage were euthanized via immersion in ice cold water for 30 minutes and collected in four 1.5 mL eppendorf tubes (30 larvae per tube for a total of 120 larvae per genotype). Larvae were washed with 1X phosphate buffered saline (PBS), PBS solution was removed, and 1 mL of Accumax (Innovative Cell Technologies Inc.; AM105) dissociation solution was added to each tube. Samples were dissociated via pipetting with a 200 µL pipette for a maximum one-hour period of time. Once the cells were fully dissociated, the sample tubes were centrifuged at 700g for five minutes. The supernatant was carefully removed, and samples resuspended in 1 mL 1X PBS, centrifuged at 700 g for five minutes, supernatant removed, and cells resuspended in 500 µL of cell medium solution (FluoroBrite DMEM (ThermoFisher; A1896701)) with 10% heat-inactivated fetal bovine serum (FBS)). Cells from the four separate tubes were then pooled together and filtered through a 70 µM strainer to remove cell aggregates. The strainer was rinsed once with 500 µL of cell medium solution. Cell viability was assessed by taking a 20 µL aliquot of each sample and staining with acridine orange/propidium iodide (Logos Biosystems; F23011) and assessing proportions of live and dead cells on a LUNA-FL™ Dual Fluorescence Cell Counter (Logos Biosystems; L20001). Samples with greater than 90% viability were used for FACS-sorting. The cells were sorted into dsRed-positive and dsRed-negative populations using a FACS ARIA III and cell viability was subsequently assessed as stated previously. Samples with greater than 85% viability and greater than 75,000 total cells were used for subsequent DNA/RNA extraction. This procedure was repeated three times to generate three biological replicates.

### Liver histology

To obtain liver tissues for histological sections and stains, zebrafish were euthanized in an ice bath for ten minutes. Liver tissues were dissected from each fish and promptly placed in 4% paraformaldehyde (PFA) and incubated overnight at 4°C with shaking. The next day, samples were washed three times in phosphate buffered saline solution (PBS) and then placed in 70% ethanol. Samples were sent to HistoServ Inc. (histoservinc.com) for sectioning of 10 µM sections and hematoxylin and eosin (H&E) staining. Slides were then imaged on a Leica DMI 6000B inverted compound microscope with a Leica DMC6200 camera, Leica Application Suite X software, and 10X 0.3 NA air, 20X 0.8 NA air, and 40X 1.25 NA oil Leica objective lenses.

### Liver LipidTox Stains

Whole livers were dissected from zebrafish after anesthetization in an ice bath for ten minutes. Livers were promptly placed in 4% PFA and rocked at room temperature for two hours. Liver tissues were washed three times in phosphate buffered saline (PBS, pH 7.2 Gibco; 20012-027) solution, and then chopped into small 10 mm pieces using fine iridectomy scissors. The PBS solution was then removed and replaced with a LipidTox solution containing 2 µL of HCS LipidTOX™ dye (ThermoFisher; H34477) in 1 mL of PBS (1:500 dilution). Samples were rocked for two hours at room temperature then washed two times with PBS. The PBS was removed and replaced with a solution containing 1:1,000 DAPI (4′,6-diamidino-2-phenylindole) in 1 mL PBS. Samples were rocked for 15 minutes in the PBS supplemented with DAPI and then washed three times with 1X PBS. LipidTox stained liver samples were then stored in the dark and imaged on a Nikon Ti2 inverted microscope with Yokogawa CSU-W1 spinning disk confocal (Hamamatsu Orca Fusion-BT camera).

### Biochemical Assays

In order to measure guanine and uric acid concentrations in zebrafish larvae, 25 *ppat* mutant and WT fish at the 5 dpf stage were euthanized and washed once in ice cold phosphate buffered saline solution (1X PBS). Larvae were spun down, supernatant removed, and then 400 µL of ice cold 1X PBS added to each tube. Larvae were homogenized until no large tissue pieces were present and then centrifuged for 10 minutes at 15,000*g* at 4°C. The clean supernatant protein extract was then transferred to a new 1.5 mL Eppendorf tube and used for protein concentration analysis and subsequent biochemical assays. If not used immediately, the protein extracts were stored at -80°C until assayed. The protein concentration of each sample was measured using the Pierce™ BCA Protein Assay Kit (23225). Each sample was then diluted to a concentration of 500 µg/mL for subsequent assays. To determine the concentration of uric acid and guanine in larva, we used the Amplex Red Uric Acid/Uricase assay kit (Molecular Probes by Life Technologies; A22181,) to measure uric acid and the Fluorometric Guanine Assay Kit (Cell Bio Labs; MET-5148) to measure guanine, following the manufacturer’s protocol. For each sample, we used 50 µL of protein extract in duplicates for the quantification assay and a standard curve with uric acid concentrations ranging from 0 µM (PBS only control) to 400 µM and guanine standard concentrations ranging from 0 µM to 200 µM concentrations. The fluorescence reading from each microplate well was measured using a Synergy H1 Microplate Reader (Biotek) using 545 nm excitation and 590 nm emission detection settings. The metabolite concentrations were calculated using the standard curve equations for each sample after normalizing to the 0 µM control.

### Statistical analyses

GraphPad Prism (version 10.1.0) and R statistical software (version 4.3.2) were used to perform statistical tests. Most statistical analyses were conducted using two-tailed unpaired Student’s t-test (for analysis of two groups) or one-way ANOVA with post-hoc Tukey’s test (for analysis of three groups) unless stated otherwise. Schematic diagrams were created using BioRender.com.

## References and Notes

1. F. N. Carelli, G. Sharma, J. Ahringer, Broad Chromatin Domains: An Important Facet ofGenome Regulation. Bioessays 39, (2017).

2. A. M. Karnay, F. Elefant, in Handbook of Epigenetics (Second edition). (Academic Press, 2017), chap. Drosophila Epigenetics, pp. 205–229.

3. D. Wenzel, F. Palladino, M. Jedrusik-Bode, Epigenetics in C. elegans: facts andchallenges. Genesis 49, 647–661 (2011).

4. J. A. Hackett et al., Promoter DNA methylation couples genome-defence mechanisms to epigenetic reprogramming in the mouse germline. Development 139, 3623–3632 (2012).

5. D. M. Maatouk et al., DNA methylation is a primary mechanism for silencing postmigratory primordial germ cell genes in both germ cell and somatic cell lineages. Development 133, 3411–3418 (2006).

6. M. E. Potok, D. A. Nix, T. J. Parnell, B. R. Cairns, Reprogramming the maternal zebrafish genome after fertilization to match the paternal methylation pattern. Cell 153,759-772 (2013).

7. Y. Stelzer, C. S. Shivalila, F. Soldner, S. Markoulaki, R. Jaenisch, Tracing dynamicchanges of DNA methylation at single-cell resolution. Cell 163, 218–229 (2015).

8. T. Matsuda, C. L. Cepko, Controlled expression of transgenes introduced by in vivo electroporation. Proc Natl Acad Sci U S A 104, 1027–1032 (2007).

9. L. Schermelleh et al., Dynamics of Dnmt1 interaction with the replication machinery andits role in postreplicative maintenance of DNA methylation. Nucleic Acids Res 35, 4301–4312 (2007).

10. M. Farooq et al., Histone deacetylase 3 (hdac3) is specifically required for liver development in zebrafish. Dev Biol 317, 336–353 (2008).

11. H. Xu, F. Wang, H. R. Kranzler, J. Gelernter, H. Zhang, Alcohol and nicotine codependence-associated DNA methylation changes in promoter regions of addiction-related genes. Sci Rep 7, 41816 (2017).

12. D. M. McCarthy et al., Transgenerational transmission of behavioral phenotypes produced by exposure of male mice to saccharin and nicotine. Sci Rep 10, 11974 (2020).

13. S. Feng, S. E. Jacobsen, W. Reik, Epigenetic reprogramming in plant and animaldevelopment. Science 330, 622–627 (2010).

14. M. A. Dawson, The cancer epigenome: Concepts, challenges, and therapeuticopportunities. Science 355, 1147–1152 (2017).

15. T. Yadav, J. P. Quivy, G. Almouzni, Chromatin plasticity: A versatile landscape thatunderlies cell fate and identity. Science 361, 1332–1336 (2018).

16. M. Gemberling, T. J. Bailey, D. R. Hyde, K. D. Poss, The zebrafish as a model forcomplex tissue regeneration. Trends Genet 29, 611–620 (2013).

17. R. Ben-Yair et al., H3K27me3-mediated silencing of structural genes is required for zebrafish heart regeneration. Development 146, (2019).

18. G. Waldo et al., Rapid protein-folding assay using green fluorescent protein. Nat Biotechnol 17, 691–695 (1999).

19. B. G. Reid, G. C. Flynn, Chromophore formation in green fluorescent protein. Biochem 36, 6786–6791 (1997).

20. K. Takayama, N. Shimoda, S. Takanaga, S. Hozumi, Y. Kikuchi, Expression patterns of dnmt3aa, dntm3ab, and dnmt4 during development and fin regeneration in zebrafish. Gene Expr Patterns 14 105-110 (2014).

21. Lee, H.J., Hou, Y., Chen, Y. et al. Regenerating zebrafish fin epigenome is characterized by stable lineage-specific DNA methylation and dynamic chromatin accessibility. Genome Biol 21, 52 (2020).

22. T. G. Pipalia et al., Cellular dynamics of regeneration reveals role of two distinct Pax7stem cell populations in larval zebrafish muscle repair. Dis Model Mech 9, 671–684 (2016).

23. J.B. Spofford, Single-locus modification of position-effect variegation in Drosophila melanogaster. O. White variegation. Genetics 57, 751–766 (1967).

24. S. C. R. Elgin, G. Reuter, Position-effect variegation, heterochromatin formation, and gene silencing in Drosophila. Cold Spring Harb. Perspec. Biol. 5 (2013).

25. A. Ashe et al., piRNAs can trigger a multigenerational epigenetic memory in the germline of C. elegans. Cell 150, 88–99 (2012).

26. A. C. Miller, N. D. Obholzer, A. N. Shah, S. G. Megason, C. B. Moens, RNA-seq-based mapping and candidate identification of mutations from forward genetic screens. Genome Res 23, 679–686 (2013).

27. I. Leshchiner et al., Mutation mapping and identification by whole-genome sequencing. Genome Res 22, 1541–1548 (2012).

28. Y. Liu, M. Chen, Histone methylation regulation as a potential target for non-alcoholic fatty liver disease. Curr Protein Pept Sci 24, 465–476 (2023).

29. Y. Chu et al., Liver Med23 ablation improves glucose and lipid metabolism through modulating FOXO1 activity. Cell Res 24, 1250-1265 (2014).

30. G. Chen et al., CYP2J2 overexpression attenuates nonalcoholic fatty liver disease induces by high-fat diet in mice. Am J Physiol Endocrinol 308, E97–E110 (2015).

