## Supplemental Data Part 1 for "A Novel Transgenic Reporter to Study Vertebrate Epigenetics"

### Supplementary Figures

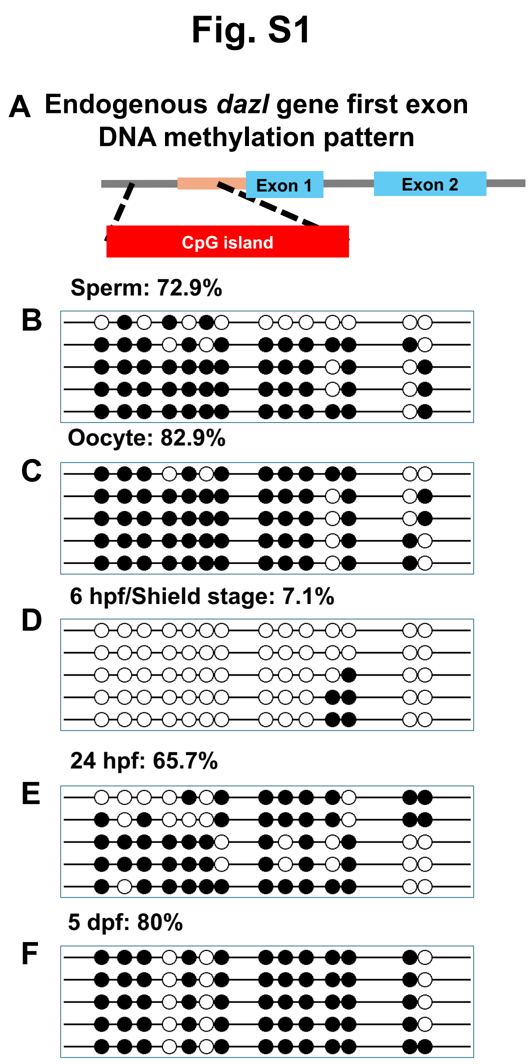


**Fig. S1. Targeted bisulfite sequencing of the endogenous *dazl* CpG island.**

(**A**) Schematic diagram showing position of the *dazl* CpG island upstream of the first exon tested for DNA methylation at different developmental stages and mature gametes. (**B-F**) bisulfite sequencing reads of *dazl* CpG island from the genomic DNA of mature sperm (**B**), mature oocytes (**C**), 6 hpf embryos (**D**), 24 hpf (**E**) and 5 dpf (**G**). Open circles represent unmethylated cytosine residues and filled circles represent methylated cytosine residues from CpG dinucleotides.

### Fig. S2

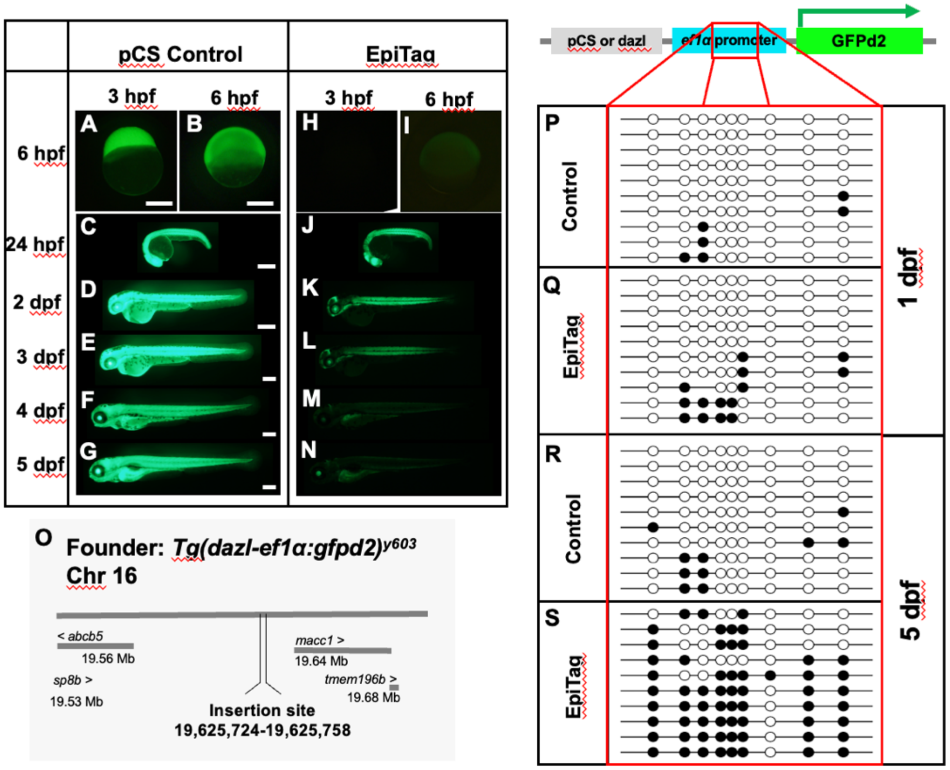


**Fig. S2. Dynamic GFPd2 expression and methylation changes of the transgene insertion during embryonic and larval development.**

**(A-G)** Green epifluorescence images of control-pCS, *Tg(pcs-ef1α:gfpd2)*$^{y605}$ and **(H-N)** of EpiTag, *Tg(dazl-ef1α:gfpd2)*$^{y603}$ embryonic and larval development. (**A,H**) 4 hpf, (**B,I**) 6 hpf, (**C,J**) 24 hpf, (**D,K**) 2 dpf, (**E,L**) 3 dpf, (**F,M**) 4 dpf and (**G,N**) 5 dpf. Scale bars = 200□m. (**O**) The EpiTag founder was isolated and mapped as described in the methods. Schematic diagram of the transgene and position of the *ef1α* promoter in the transgene. Red box noting approximate region of the *ef1α* promoter that was assayed for change in methylation. (**P-S**) Representative bisulfite sequencing reads of genomic DNA from 1 dpf embryos (**P,Q**) and 5 dpf larvae (**R,S**). Open circles represent unmethylated cytosine residues and filled circles represent methylated cytosine residues from CpG dinucleotides.

#### Fig. S3

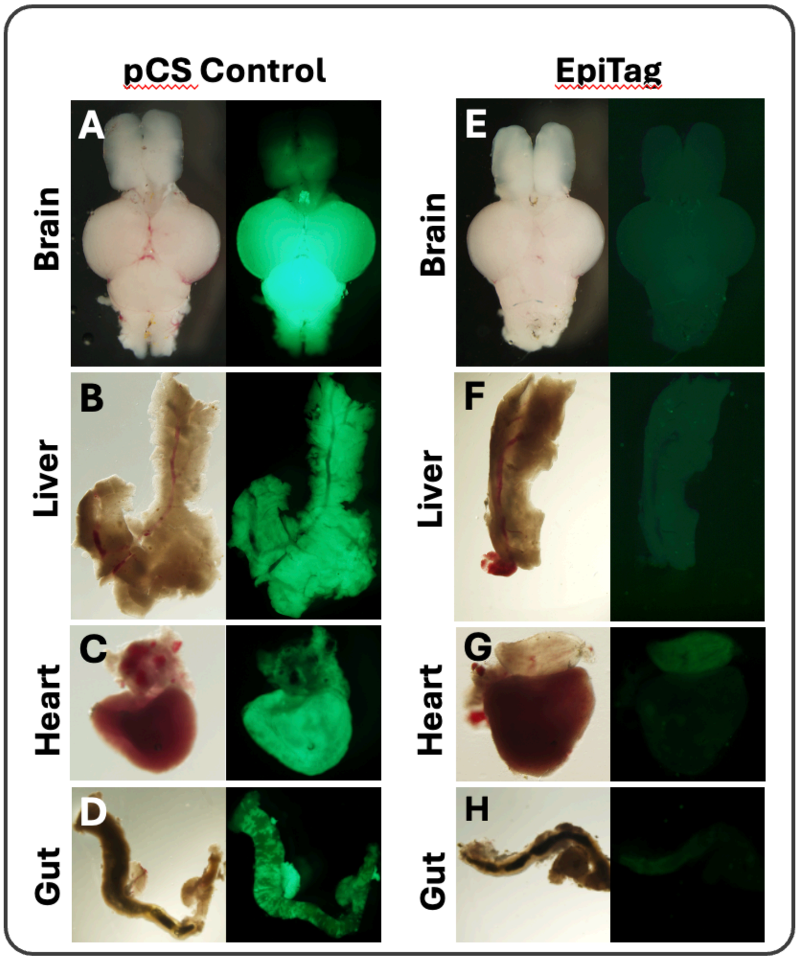


**Fig. S3. EpiTag line silencing in internal adult organs.**

(**A-H**) Paired bright field (left) and green epifluorescence (right) images of dissected internal organs from pCS Control (**A-D**) and EpiTag (**E-H**) transgenic animals. Brain (**A,E**), liver (**B,F**), heart (**C,G**) and gut (**D,H**) are shown. PCS control and EpiTag images were collected using the same settings for comparison.

### Fig. S4

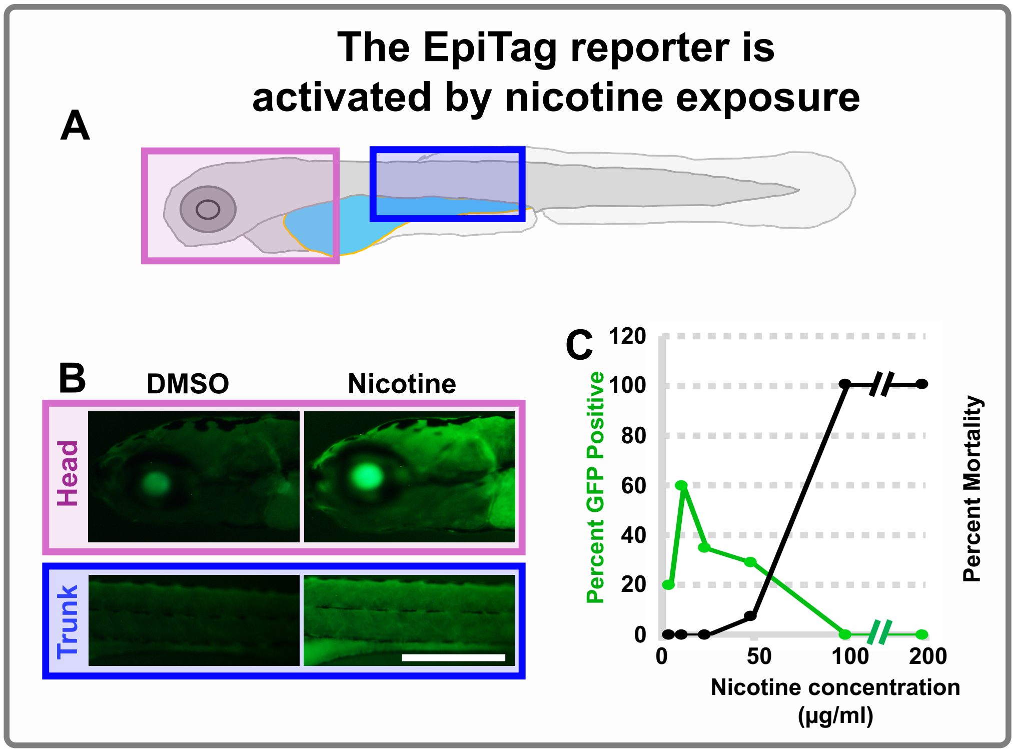


**Fig. S4. Treatment of EpiTag larvae with nicotine results in GFPd2 activation at levels below those that induce mortality.**

**(A)** Schematic diagram of a 5 dpf EpiTag larvae with magenta and blue boxes noting the approximate regions of the head and trunk imaged in panel B. (**B**) Green epifluorescence images of the heads (top) or trunks (bottom) of 5 dpf larvae treated with either DMSO (left) or nicotine (right). (**C**) Quantification of the percentage of GFP positive (green) or severely malformed/dead (black) 5 dpf EpiTag larvae after treatment from 3-5 dpf with the indicated concentrations of nicotine. Scale bar, 200 µm.

### Fig. S5

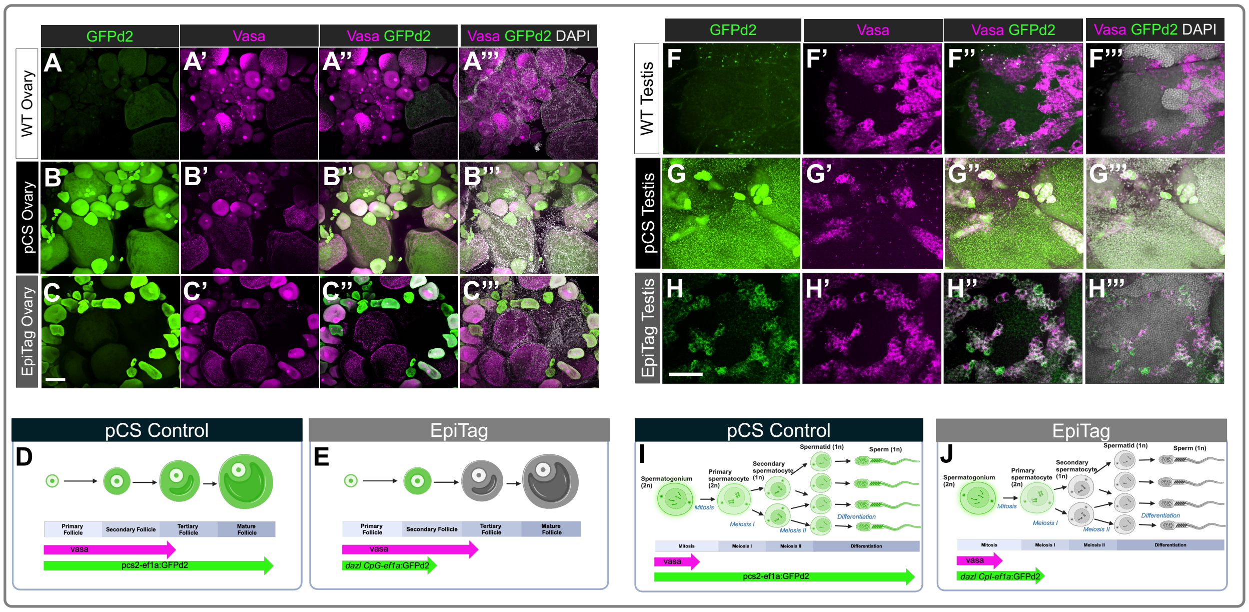


**Fig. S5. GFPd2 expression in early stage oocytes and sperm colocalizes with Vasa**

(**A-C**) Whole-mount ovary tissues of wild-type (**A**), *Tg(pcs-ef1α:gfpd2)*$^{y605}$ Control (**B**), and *Tg(dazl-ef1α:gfpd2)*$^{y603}$ EpiTag (**C**) adult females stained with *gfpd2* and *vasa* HCR probes and DAPI nuclear dye. Scale bar, 25 µm. (**D,E**) Schematic diagram showing different stages of oogenesis starting from the primary follicle and the corresponding GFPd2 expression patterns in pCS control (**D**) and EpiTag (**E**) fish. (**F-H**) Whole-mount testes tissues of wild-type (**F**), *Tg(pcs-ef1α:gfpd2)*$^{y605}$ Control (**G**), and *Tg(dazl-ef1α:gfpd2)*$^{y603}$ EpiTag (**H**) adult males stained with *gfpd2* and *vasa* HCR probes and DAPI nuclear dye. Scale bar, 25 µm (**I,J**) Schematic diagram showing different stages of spermatogenesis starting from the spermatogonium stage and the corresponding GFPd2 expression patterns in pCS Control (**I**) and EpiTag (**J**) fish.

#### Fig. S6

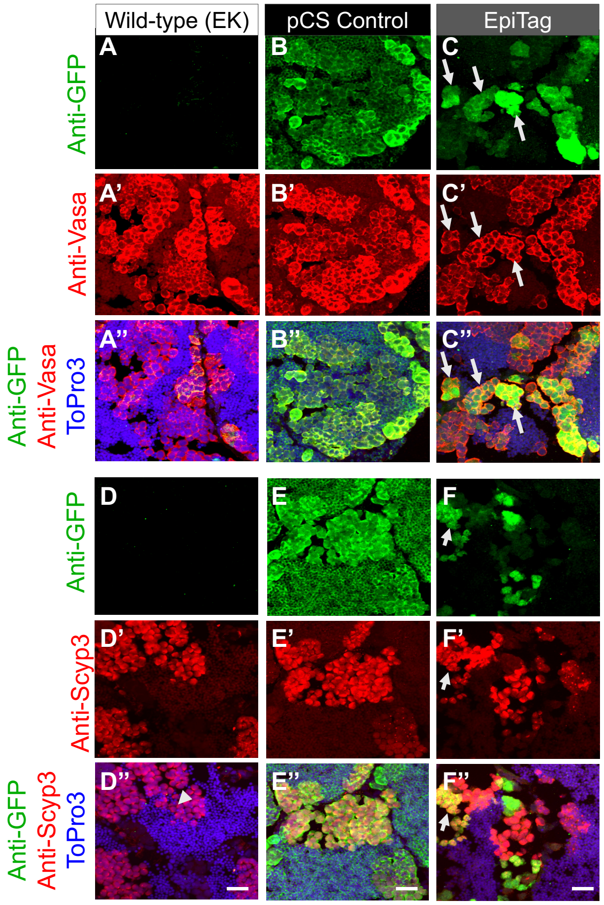


**Fig. S6. GFPd2 reactivation during spermatogenesis**

(**A-C''**) Immunohistochemical staining of cryosections from zebrafish testes probed for Anti-GFP (green) and anti-Vasa (red) and co-stained with ToPro3 nuclear dye (blue). (**D-F''**) Immunohistochemistry of cryosections from zebrafish testes probed for Anti-GFP (green) and anti-Scyp3 (red) and co-stained with ToPro3 nuclear dye (blue). Images shown are sections from wild-type (**A-A'', D-D''**), pCS control (**B-B'', E-E"**) and EpiTag (**C-C", F-F"**) zebrafish testes. Scale bar, 20 μm.

### Fig. S7

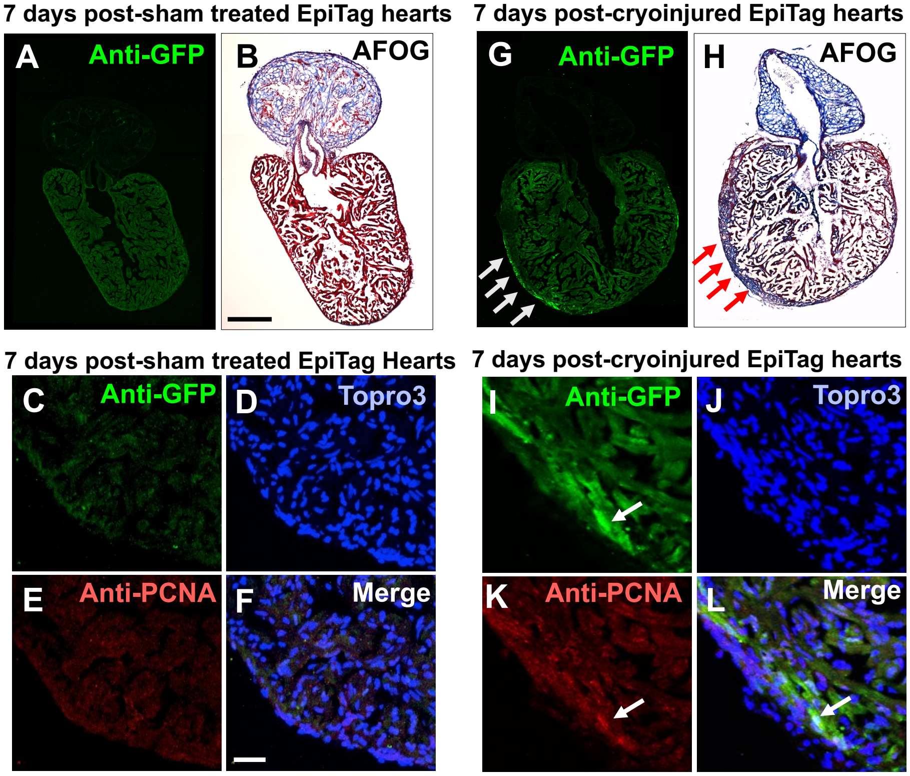


**Fig. S7. GFPd2 reactivation during heart regeneration.**

Immunohistochemical staining of cryosections of EpiTag zebrafish hearts either seven days post-sham treatment (**A-F**) or seven days post-cryoinjury (**G-L**). (**A,B**) and (**G,H**) are serial sections and panels (**C-F**) and panels (**I-L**) are showing the same sections. Anti-GFP (green) is shown in panels (**A,C,G,I**), anti-PCNA (red) is shown in panels (**E,K**), nuclei stained with ToPro3 (blue) are shown in panels (**D,J**) and merged images are shown in panels (**F,L**). AFOG staining is shown in panels (**B,H**). Scale bars = 200 µm (**A,B,G,H**) and 20 µm (**C-F,I-L**).
