## Supplemental Data Part 2 for "A Novel Transgenic Reporter to Study Vertebrate Epigenetics"

#### Fig. S8

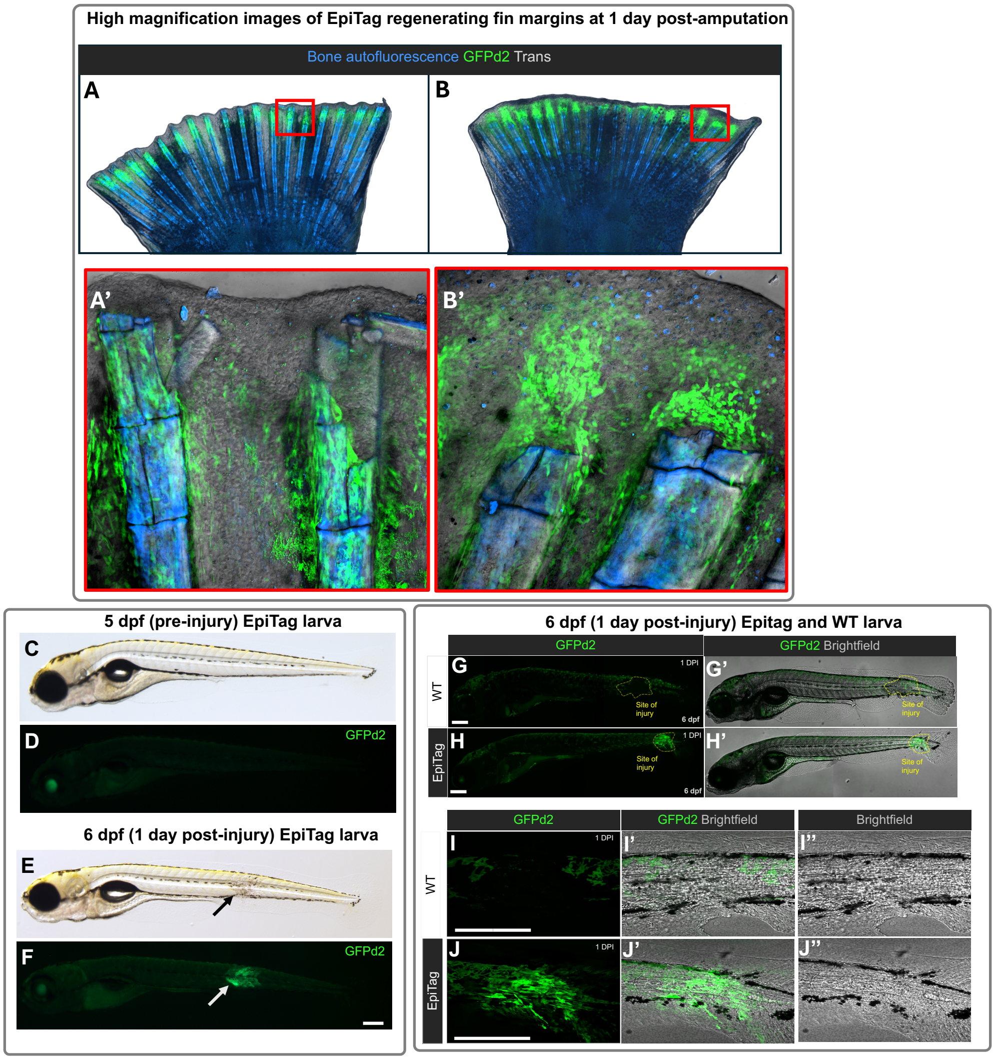


**Fig. S8. GFPd2 reactivation during fin and somite regeneration.**

(**A,B**) Confocal images of 1 day post-amputation recovering EpiTag tail fin. Red boxes in A and B note the approximate areas of the regenerating fin imaged at higher magnification (**A',B'**). GFPd2 expressing cells are in green and autofluorescence from osteoblasts is in blue. (**C-F**) Bright field (**C,E**) and green epifluorescence images (**D,F**) of the same EpiTag larva at 5 dpf, just prior to trunk cryoinjury (**C,D**), and one day later at 6 dpf, one day post-injury (dpi) (**E,F**). Scale bar, 50µm. (**G,H**) Representative GFP confocal and brightfield images (**G',H'**) of 6 days post fertilization (6 dpf) wild-type (**G**) and EpiTag (**H**) larvae 1 day post somite injury (dpi). Scale bar, 200 µm. (**I,J**) Zoomed in GFP (**I,J**) and brightfield (**I'',J''**) confocal images of the injury site of wild-type (**I**) and EpiTag (**J**) larvae 1 dpi. Scale bar, 200 µm.

### Fig. S9

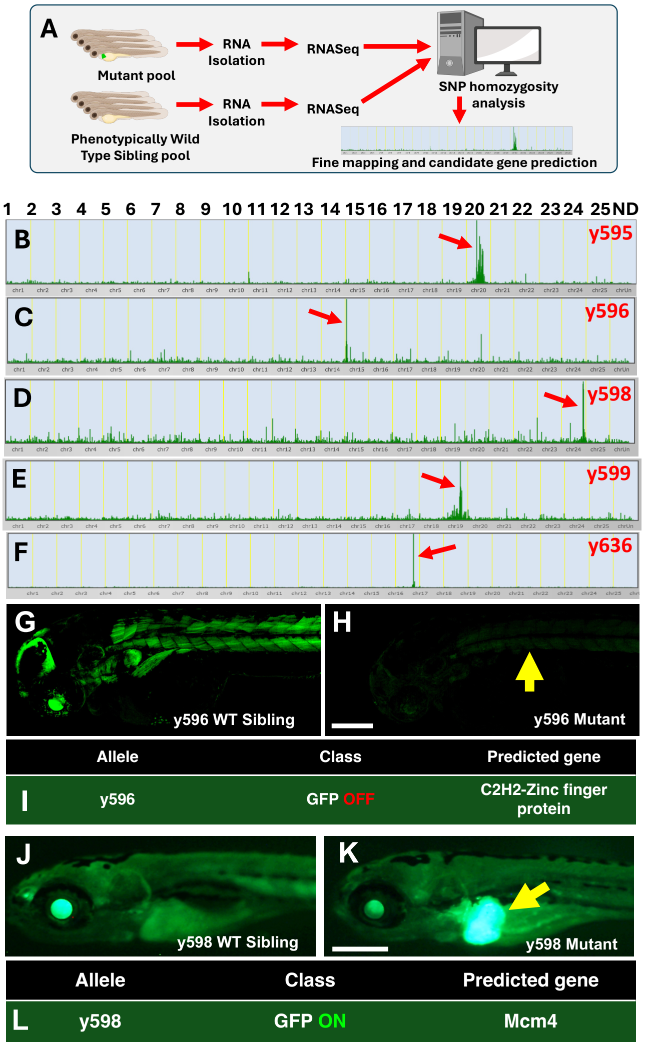


**Fig. S9. RNAseq based mapping of epigenetic mutants.**

**(A)** Schematic diagram showing protocol of mutant mapping using RNAseq based approach. **(B-F)** Homozygosity plots showing mapping results of different mutants (red arrows), (**B**) *y595*, (**C**) *y596*, (**D**) *y598*, (**E**) *y599*, and (**F**) *y636.* Lateral view green epifluorescence images of a y596 WT EpiTag fish (**G**) and its y596 mutant EpiTag sibling (**H**) at 2 dpf, showing ubiquitously decreased GFPd2 expression (arrow). Scale bar, 250 µm. (**I**) Predicted mutant allele of the ubiquitously silenced y596 mutant. Lateral view green epifluorescence images of a y598 WT EpiTag fish (**J**) and its y598 mutant EpiTag sibling (**K**) at 5 dpf, showing strongly increased GFPd2 expression in the liver (arrow). Scale bar, 250µm. (**L**) Predicted mutant allele of the "liver GFP on" y598 mutant.

#### Fig. S10

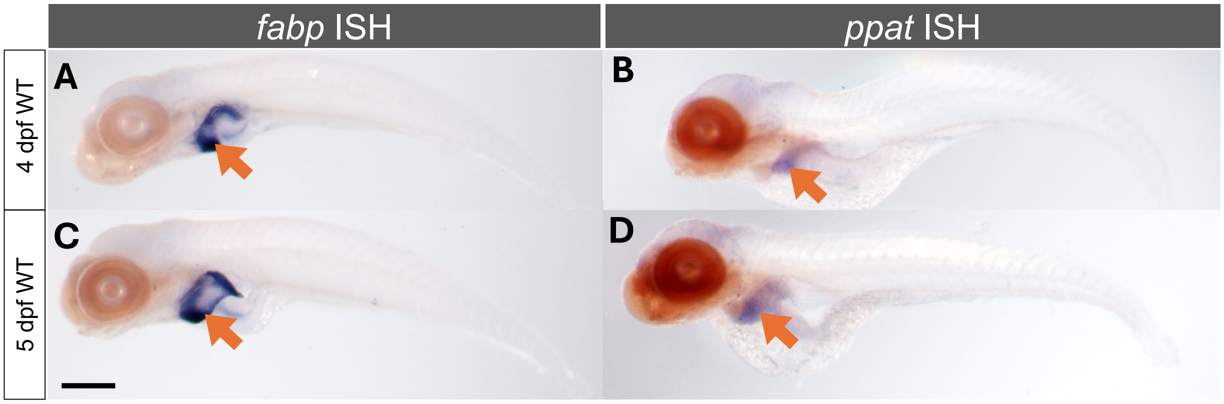


**Fig. S10.** ***Ppat*** **is expressed in liver tissues of 4 dpf and 5 dpf zebrafish larvae**

(**A,B**) 4 dpf wild-type fish stained with a liver-specific riboprobe *fabp10a* (**A**) and a *ppat* riboprobe (B) with the livers noted with an orange arrow. (**C,D**) 5 dpf wild-type fish stained with a liver-specific riboprobe *fabp10a* (**C**) and a *ppat* riboprobe (**D**) with the livers noted with an orange arrow. Scale bar, 200 µm.

### Fig. S11

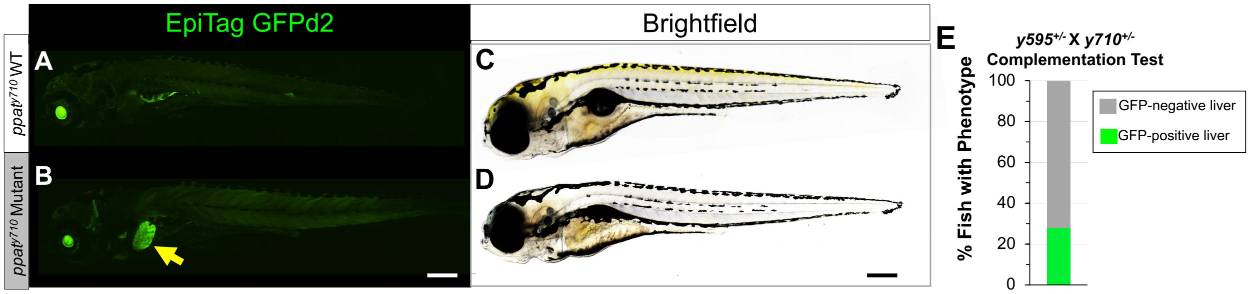


**Fig. S11.** ***ppat*$^{y710}$ CRISPR mutants phenocopy *y595* mutants and the two mutants fail to complement one another**

**(A,B)** Representative GPFd2 epifluorescence images of a *ppat*$^{y710}$ WT larvae (**A**) and a *ppat*$^{y710}$ mutant sibling (**B**) at 5 dpf, with liver-specific GFPd2 activation noted with a yellow arrow, and the corresponding brightfield images of the *ppat*$^{y710}$ embryo (C) and its *ppat*$^{y710}$ mutant counterpart (**D**). (**E**) Complementation test quantification of the percentage of fish progeny from a y710 and y595 heterozygous cross exhibiting GFPd2 activation in liver tissues.

#### Fig. S12

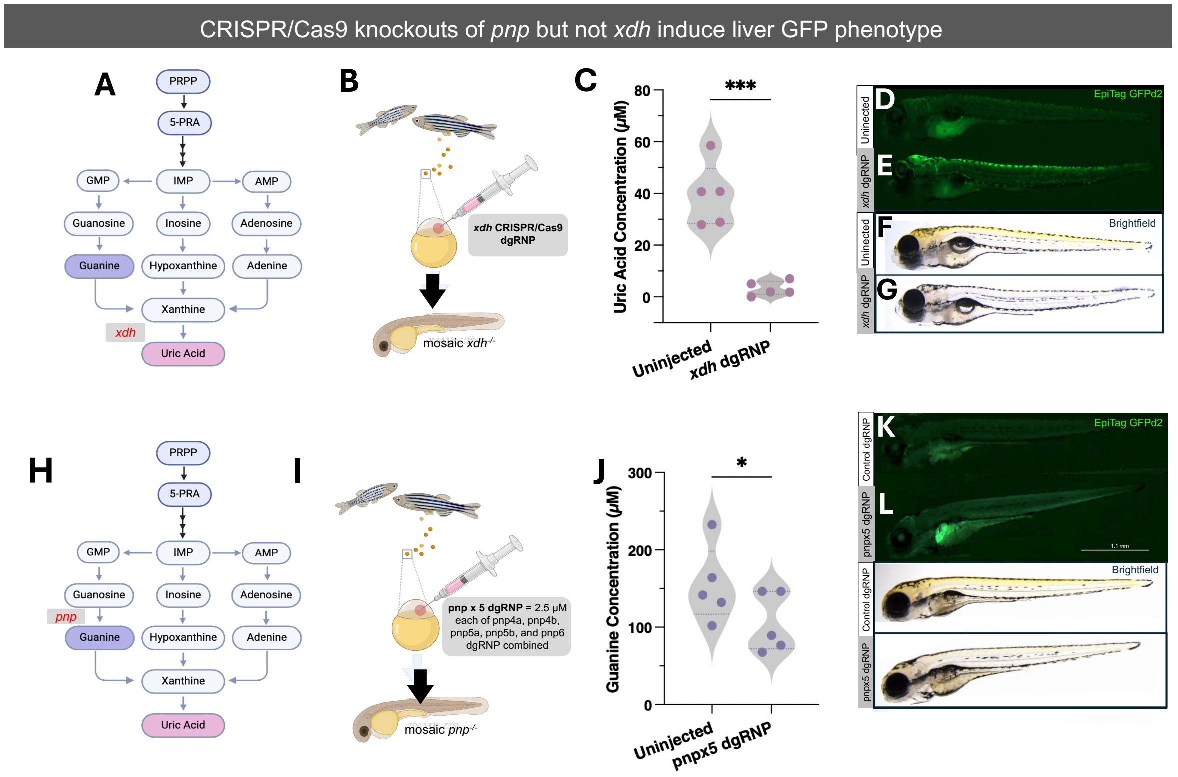


**Fig. S12. Guanine, but not uric acid, depletion results in GFPd2 activation in EpiTag fish liver tissues.**

(**A**) *De novo* purine synthesis pathway with the enzymatic step of *xdh* noted in red. (**B**) Schematic diagram of the CRISPR/Cas9 dgRNP injection strategy to generate mosaic F0 *xdh* mutants. (**C**) Quantification of uric acid concentrations in 5 dpf uninjected or *xdh* dgRNP injected sibling fish. (**D-E**) Representative green epifluorescence images of 5 dpf EpiTag control fish (**D**) and an *xdh* dgRNP injected sibling (**E**). Brightfield image of the 5 dpf EpiTag control fish (**F**) and an *xdh* dgRNP injected sibling (**G**) with a lack of xanthophore pigmentation. (**H**) *De novo* purine synthesis pathway with the enzymatic step of *pnp* noted in red. (**I**) Schematic diagram of the CRISPR/Cas9 dgRNP injection strategy to generate mosaic F0 *pnp* mutants via targeting of all five *pnp* genes. (**J**) Quantification of guanine concentrations in 5 dpf uninjected or *pnp* dgRNP injected sibling fish. (**K-L**) Representative green epifluorescence images of 5 dpf EpiTag control fish injected with a control dgRNP (**K**) and a *pnp* dgRNP injected sibling (**L**). Brightfield image of the 5 dpf EpiTag control fish (**M**) and a *pnp* dgRNP injected sibling (**N**). Scale bar, 200 µm. For (C, J): n=5 pooled replicates, $^*P < 0.05$, $^{***}P < 0.001$; as determined by paired, two-tailed Student's *t* test.

#### Fig. S13

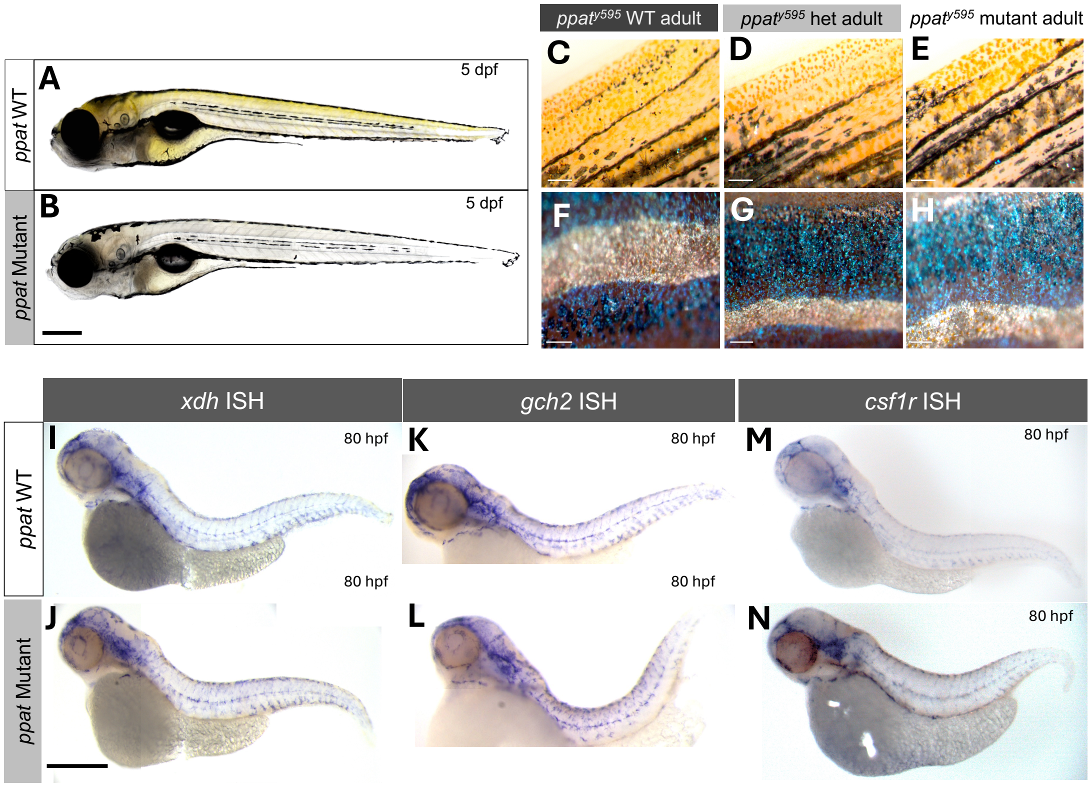


**Fig. S13. *Ppat* mutants show defects in xanthophore pigmentation in larval, but not adult, stages and exhibit normal xanthophore cell development.**

**(A)** Brightfield image of a 5 dpf *ppat* WT fish with normal yellow xanthophore pigmentation and its *ppat* mutant sibling (**B**) which lacks xanthophore pigmentation at this stage/ Scale bar, 200 µm. **(C-H)** Images of WT **(C,F),** heterozygote **(D,G),** and mutant **(E,H)** adult fins **(C-E)** and trunk **(F-H)** showing normal xanthophore, iridophore, and melanophore pigment cell presence and pigmentation. *In situ* hybridization (ISH) images of xanthophore cell marker genes *xdh* (**I, J**), *gch2* (**K, L**), and *csf1r* (**M, N**) in 80 hpf WT (**I, K, M**) and *ppat* mutant (**J, L, N**) fish. Scale bar, 200 µm.

### Fig. S14

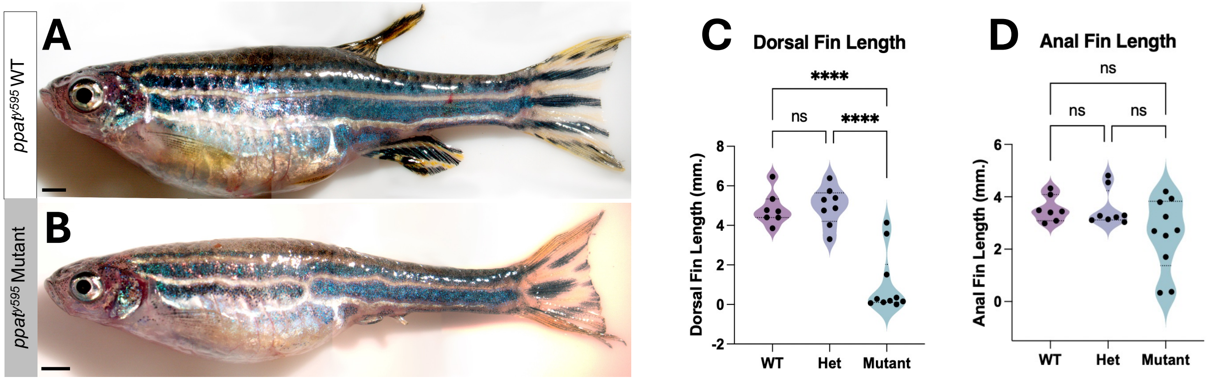


**Fig. S14. *Ppat* mutants exhibit abnormal medial fin development**

Representative image of 4-month old *ppat*$^{y595}$ WT adult (**A**) and mutant sibling with abnormal dorsal and anal fin morphology (**B**). **(C,D)** Quantification of *ppat*$^{y595}$ WT and mutant dorsal (**C**) and anal fin (**D**) lengths at 4-months of age. For (**C, D**): n=10, ****P < 0.0001; as determined by unpaired, two-tailed Student's *t* test.

#### Fig. S15

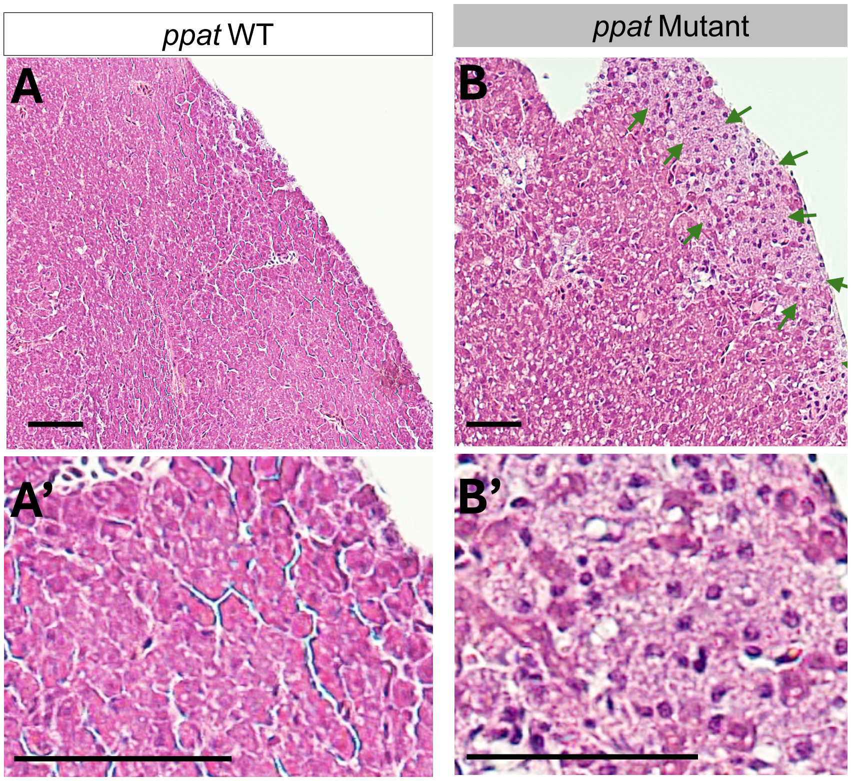


**Fig. S15.** ***Ppat*** **mutant liver tissues exhibit signs of fatty liver disease.**

(**A**) 20X magnification image of a WT liver tissue displaying normal hepatocyte morphology. (**B**) 20X magnification image of a *ppat*$^{y595}$ mutant liver tissue exhibiting ballooned hepatocytes (green arrows). (**A', B'**) Zoomed in images of A and B. Scale bar, 50 µm.

#### Fig. S16

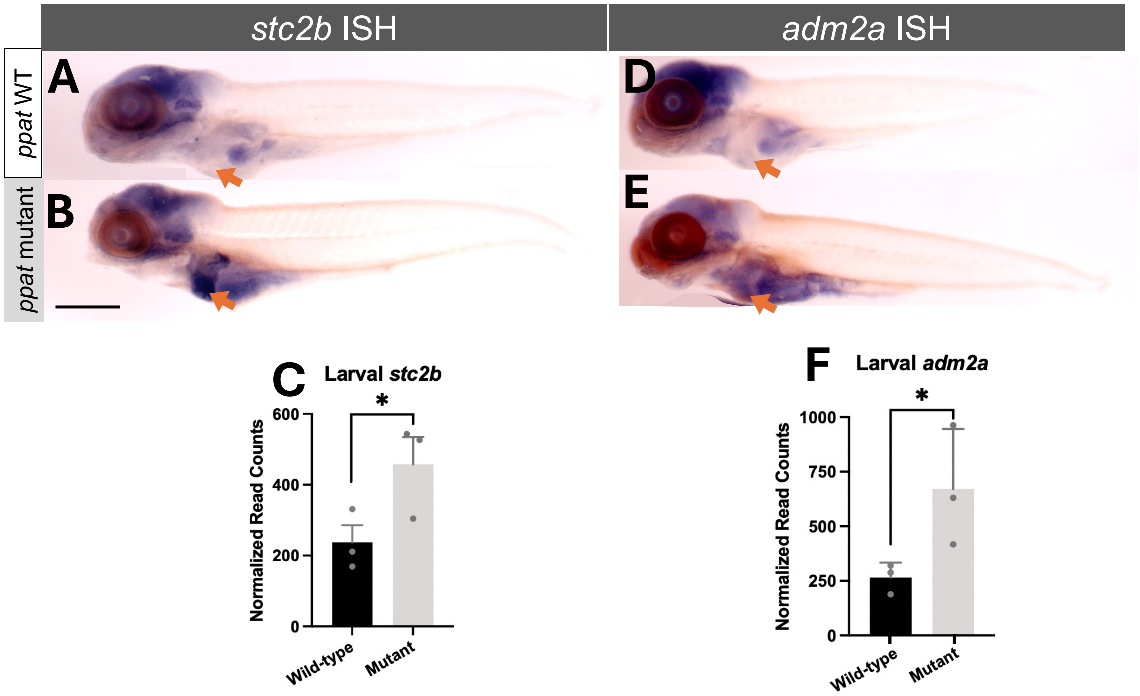


**Fig. S16. *In situ* hybridization of upregulated genes in *ppat* mutant larval and adult liver tissues**

*In situ* hybridization of *stc2b* in 6 dpf WT (**A**) and *ppat* mutant (**B**) larvae, with the liver tissues highlighted (orange arrow). (**C**) Normalized read counts of *stc2b* in wild-type and *ppat* mutant 6 dpf larval liver tissues. *In situ* hybridization of *adm2a* in 6 dpf WT (**D**) and *ppat* mutant (**E**) larvae, with the liver tissues highlighted (orange arrow). (**F**) Normalized read counts of *adm2a* in wild-type and *ppat* mutant 6 dpf larval liver tissues.

#### Fig. S17

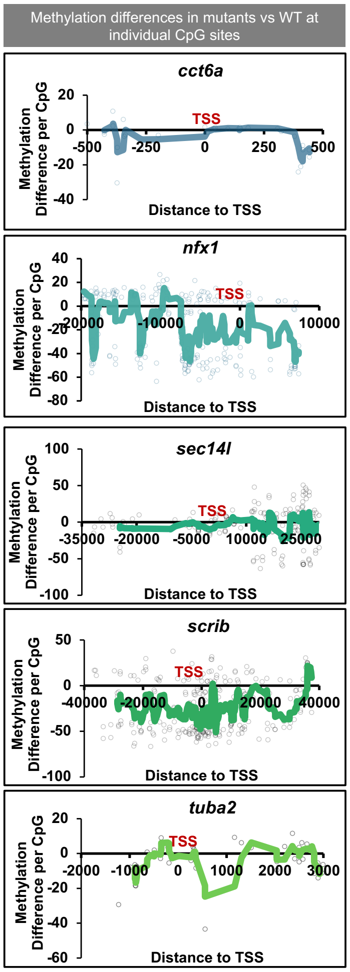


**Fig. S17. Promotor and gene body hypomethylation of genes associated with MAFLD**

Graphs displaying the adult *ppat* mutant average methylation difference at individual CpG sites surrounding the promoter and gene body region of genes that are associated with metabolic dysfunction-associated fatty liver disease (MAFLD).

**Fig. S18**

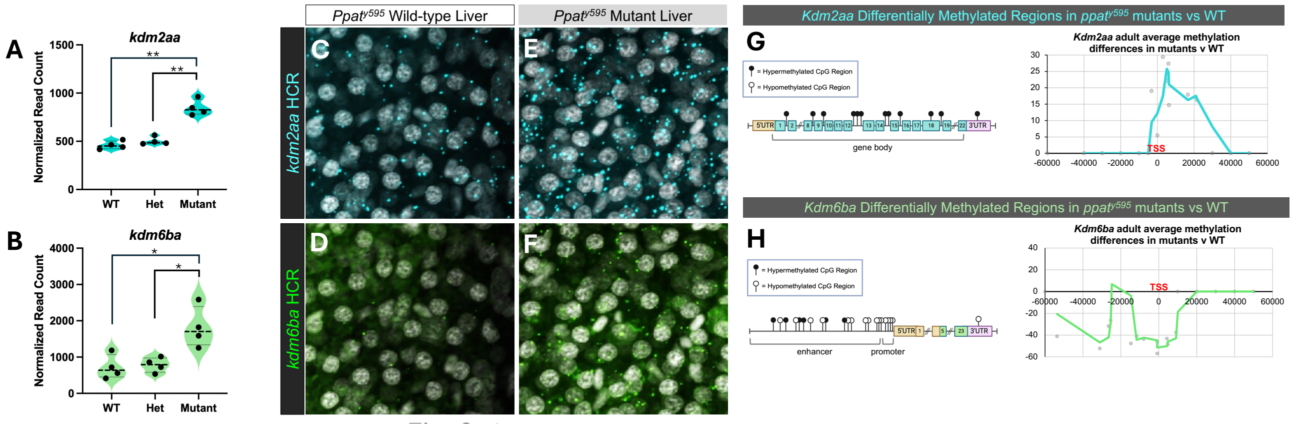


**Fig. S18. Histone demethylase genes *kdm2aa* and *kdm6ba* display differential expression and methylation in *ppat* mutant livers**

Normalized read counts of *kdm2aa* (**A**) and *kdm6ba* (**B**) from *ppat*$^{y595}$ WT, heterozygote, and mutant adult livers. (**C-F**) Whole-mount *in situ* hybridization chain reaction (HCR) of *kdm2aa* (**C**) and *kdm6ba* (**D**) in wild-type liver tissues and of *kdm2aa* (**E**) and *kdm6ba* (**F**) in *ppat* mutant liver tissues co-stained with DAPI, demonstrating increased transcript abundance in mutant hepatocytes. (G) Representation of individual CpG site methylation differences

#### Fig. S19

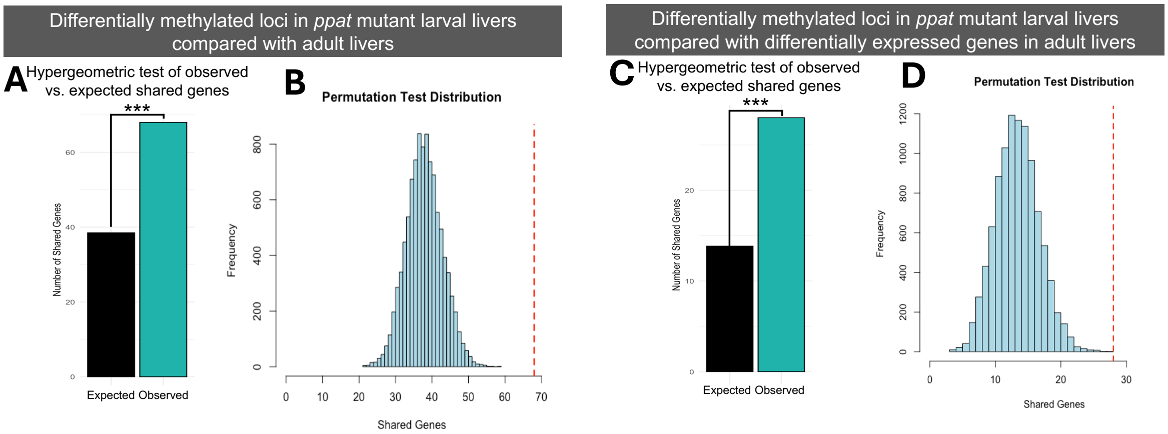


**Fig. S19. Statistical tests comparing observed vs. expected shared loci between different datasets**

**(A)** Hypergeometric test of expected number of shared loci between larval and adult differentially methylated datasets, where the observed number of shared loci is extremely significantly higher than statistically expected (*p=1.171631E-08*). (**B**) Permutation test displaying expected distribution of shared genes between larval and adult differentially methylated loci datasets, with the observed number of shared genes labelled (dotted red line). (**C**) Hypergeometric test of expected number of shared loci between larval differentially methylated loci and adult differentially expressed loci, where the observed number of shared loci is extremely significantly higher than statistically expected (*p=8.68803E-05*). (**D**) Permutation test displaying expected distribution of shared genes between larval differentially methylated loci and adult differentially expressed loci datasets, with the observed number of shared genes labelled (dotted red line).

### Fig. S20

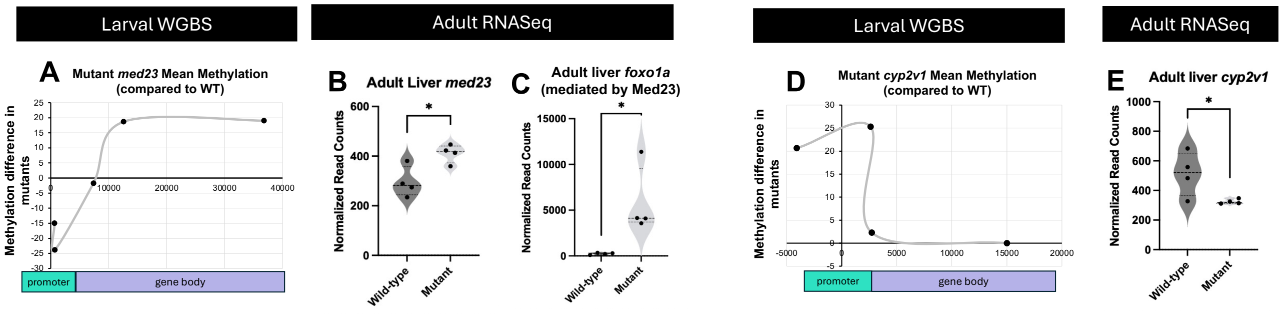


**Fig. S20. Several genes in *ppat* mutant livers exhibit early methylation changes and later expression differences**

(**A**) Average methylation differences of CpG regions of the *med23* loci in *ppat* mutant larvae compared with WT. Normalized read counts of *med23* (**B**) *and foxo1a* (**C**) in *ppat* mutant and WT adult livers. (**D**) Average methylation differences of CpG regions of the *cyp2v1* loci in *ppat* mutant larvae compared with WT. (**E**) Normalized read counts of *cyp2v1* in *ppat* mutant and WT adult livers.

**Table S1.**

| **Allele number** | **Phenotype** | **Mapped Chromosome** | **Candidate gene** | **Molecular Function** |
|---|---|---|---|---|
| ***y595*** | GFP expression in liver at 5 dpf | 20 | Ppat | Purine metabolism |
| ***y596*** | Overall weak GFP expression at 2 dpf | 15 | Novel C2H2-zinc finger protein | Binds methylated DNA |
| ***y598*** | GFP expression in liver at 5 dpf | 24 | Mcm4 | Chromatin binding, DNA replication licensing factor |
| ***y599*** | GFP expression in the pharynx at 5 dpf | 19 | ATP6V1c1b | Intracellular ATPase |
| ***y601*** | GFP expression in the blood cells at the CHT region at 5 dpf | ND | ND | |
| ***y636*** | GFP expression in the ventral side of the brain at 5 dpf | 17 | Eprs | tRNA aminoacylation for protein translation |
| ***y637*** | No GFP in the head at 2 dpf | ND | ND | |

ND: not determined

**Table S2.**

| Primer name | Primer sequence (5' to 3') | Comments |
|---|---|---|
| 5'E_dazl_F | GGGGACAACTTTGTATAGAAAAGTTGCAT AAATATTCGTCCAATTAAGATT | 5' primer to clone *dazl* CpG island into Gateway vector |
| 5'E_dazl_R | GGGGACTGCTTTTTTGTACAAACTTGCAAA AACCGAATAGTAGTTTAACAT | 3' primer to clone *dazl* CpG island into Gateway vector |
| 5'E_pcs_F | GGGGACAACTTTGTATAGAAAAGTTGGAA GGCCTCTTCGCTATTACGCCAG | 5' primer to clone pCS vector backbone sequence into Gateway vector |
| 5'E_pcs_R | GGGGACTGCTTTTTTGTACAAACTTGGTCA ATGGGGATGTACTTGGCAGCC | 3' primer to clone pCS vector backbone sequence into Gateway vector |
| ME_ef1a_F | GGGGACAAGTTTGTACAAAAAAGCAGGCT GGGCCCTCGAGCAGGGGGATCATCT | 5' primer to clone *Xenopus ef1a* promoter sequence Gateway vector |
| ME_ef1a_R | GGGGACCACTTTGTACAAGAAAGCTGGGT GCCTGCAGGAAGCTTCAGCTAGAACT | 3' primer to clone *Xenopus ef1a* promoter sequence Gateway vector |
| 3'E_GFPd2_F | GGGGACCACTTTGTACAAGAAAGCTGGGT ATGGTGAGCAAGGGCGAGGAGCTGT | 5' primer to clone GFPd2 into Gateway vector |
| 3'E_GFPd2_R | GGGGACAACTTTGTATAATAAAGTTGCTA CACATTGATCCTAGCAGAAGCAC | 3' primer to clone GFPd2 into Gateway vector |
| ppat_CRISPR_F | TGTAAAACGACGGCCAGTACTTTTGGATCAC TGCTCGATT | *ppat* forward primer to screen indels by fluorescent PCR |
| ppat_CRISPR_R | GTGTCTTCAAATTCTCTGCATTTCCACCT | *ppat* reverse primer to screen indels by fluorescent PCR |
| dazl_BS_F | TGTTTAGTTTGTTTAATGAAGAAATA | forward primer for endogenous *dazl* CpG targeted bisulfite sequencing |
| dazl_BS_R | AAACCCCCTAAACCATTTTAAATAA | Reverse primer for endogenous *dazl* CpG targeted bisulfite sequencing |
| Xef1a_BS_F | TTGAAATTTTTAGGTATGTAAGTTAGTTTA | forward primer for *Xenopus ef1a* promoter targeted bisulfite sequencing |
| Xef1a_BS_R | CAAATCCAAAATTCCCAAAATACTA | reverse primer for *Xenopus ef1a* promoter targeted bisulfite sequencing |
| GFPd2_qpcr_F | ATGCCTGCTTGCCGAATA | forward primer for *gfpd2* RT-qPCR |
| GFPd2_qpcr_R | GCCAACGCTATGTCCTGATA | reverse primer for *gfpd2* RT-qPCR |
| Xef1a_399_F | TTAATCCCCGCCCAGTAGAG | forward primer for *Xenopus ef1a* promoter qPCR |
| Xef1a_494_R | AGCTTCAGCTAGAACTCGCC | reverse primer for *Xenopus ef1a* promoter qPCR |
| ppat_ISH_F | CTCTAGCCAGCGTGGAAACA | forward primer to generate *ppat in situ* riboprobe |

| | | |
|---|---|---|
| ppat_ISH_T3-R | GAGattaaccctcactaaagggaTCAGATGAAACCACCCAGCC | reverse primer to generate *ppat in situ* riboprobe with T3 polymerase binding site |
| xdh_ISH_F | AGCCTTCAAGCAATCCCCTC | forward primer to generate *xdh in situ* riboprobe |
| xdh_ISH_T3-R | GAGattaaccctcactaaagggaTGGCCCCCACTATATGTCCA | reverse primer to generate *xdh in situ* riboprobe with T3 polymerase binding site |
| gch2_ISH_F | GCAATGGCTTTAGCGACCTG | forward primer to generate *gch2 in situ* riboprobe |
| gch2_ISH_T3-R | GAGattaaccctcactaaagggaTCACGGTACGGCTGTTCATC | reverse primer to generate *gch2 in situ* riboprobe with T3 polymerase binding site |
| csf1ra_ISH_F | CTCTGAGATGTTCTTCGCGCT | forward primer to generate *csf1ra in situ* riboprobe |
| csf1ra_ISH_T3-R | GAGattaaccctcactaaagggaGCTTCGTTCTGACCCGTACA | reverse primer to generate *csf1ra in situ* riboprobe with T3 polymerase binding site |
| fabp10a_ISH_F | GACGTGGCAGGTTTACGCT | forward primer to generate *fabp10a in situ* riboprobe |
| fabp10a_ISH_T3-R | GAGattaaccctcactaaagggaGGATGTGGGAGAATCGGTCA | reverse primer to generate *fabp10a in situ* riboprobe with T3 polymerase binding site |
| adm2a_ISH_F | AGGCACATGCCAGGTACAAA | forward primer to generate *adm2a in situ* riboprobe |
| adm2a_ISH_T3-R | GAGattaaccctcactaaagggaCCCAGCATGATTCAGTCGGA | reverse primer to generate *adm2a in situ* riboprobe with T3 polymerase binding site |
| stc2b_ISH_F | AGCGCTCATGACACATCACA | forward primer to generate *stc2b in situ* riboprobe |
| stc2b_ISH_T3-R | GAGattaaccctcactaaagggaCAGCCTCTGGGTGAGGAAAG | reverse primer to generate *stc2b in situ* riboprobe with T3 polymerase binding site |

#### Supplementary References


1. M. Westerfield, *The zebrafish book. A guide for laboratory use of zebrafish (Daniorerio).* (Univ. of Oregon Press, Eugene, ed. 4th, 2000).
2. C. B. Kimmel, W. W. Ballard, S. R. Kimmel, B. Ullmann, T. F. Schilling, Stages ofembryonic development of the zebrafish. *Dev Dyn* **203**, 253-310 (1995).
3. G. M. Her, C. C. Chiang, W. Y. Chen, J. L. Wu, In vivo studies of liver-type fatty acid binding protein (L-FABP) gene expression in liver of transgenic zebrafish (Danio rerio).*FEBS Lett* **538**, 125-133 (2003).
4. K. M. Kwan *et al.*, The Tol2kit: a multisite gateway-based construction kit for Tol2transposon transgenesis constructs. *Dev Dyn* **236**, 3088-3099 (2007).
5. K. M. Tabor *et al.*, Brain-wide cellular resolution imaging of Cre transgenic zebrafishlines for functional circuit-mapping. *Elife* **8**, (2019).
6. L. Solnica-Krezel, A. F. Schier, W. Driever, Efficient recovery of ENU-inducedmutations from the zebrafish germline. *Genetics* **136**, 1401-1420 (1994).
7. K. Labun *et al.*, CHOPCHOP v3: expanding the CRISPR web toolbox beyond genomeediting. *Nucleic Acids Res* **47**, W171-W174 (2019).
8. E. Y. Chen *et al.*, Enrichr: interactive and collaborative HTML5 gene list enrichment analysis tool. *BMC Bioinform* **128**, (2013).
9. M. V. Kuleshov *et al.*, Enrichr: a comprehensive gene set enrichment analysis web server 2016 update. *Nucleic Acids Res* **44**, W90-W97 (2016).
10. A. V. Gore *et al.*, Epigenetic regulation of hematopoiesis by DNA methylation. *Elife* **5**,e11813 (2016).
11. A. V. Gore *et al.*, An epigenetic mechanism for cavefish eye degeneration. *Nat Ecol Evol* **2**, 1155-1160 (2018).
12. R. Ibarra-Garcia-Padilla, A. G. A. Howard IV, E. W. Singleton, R. A. Uribe, A protocol for whole-mount immune-coupled hybridization chain reaction (WIHCR) in zebrafish embryos and larvae. *STAR Prot* **2**, 100709 (2021),
13. K. D. Poss, L. G. Wilson, M. T. Keating, Heart regeneration in zebrafish. *Science* **298**,2188-2190 (2002).
14. J. M. Gonzalez-Rosa, V. Martin, M. Peralta, M. Torres, N. Mercader, Extensive scarformation and regression during heart regeneration after cryoinjury in zebrafish. *Development* **138**, 1663-1674 (2011).
15. T. G. Pipalia *et al.*, Cellular dynamics of regeneration reveals role of two distinct Pax7stem cell populations in larval zebrafish muscle repair. *Dis Model Mech* **9**, 671-684 (2016).
